# GWAS-driven Pathway Analyses and Functional Validation Suggest *GLIS1* as a Susceptibility Gene for Mitral Valve Prolapse

**DOI:** 10.1101/433268

**Authors:** Mengyao Yu, Adrien Georges, Nathan R. Tucker, Sergiy Kyryachenko, Katelyn Toomer, Jean-Jacques Schott, Francesca N. Delling, Patrick T. Ellinor, Robert A. Levine, Susan A. Slaugenhaupt, Albert A. Hagège, Christian Dina, Xavier Jeunemaitre, David J. Milan, Russell A. Norris, Nabila Bouatia-Naji

**Affiliations:** INSERM, UMR970, Paris Cardiovascular Research Center, 75015 Paris, France; University Paris Descartes, Sorbonne Paris Cité, Faculty of Medicine, 75006 Paris, France; Cardiovascular Research Center, Cardiology Division, Massachusetts General Hospital, Harvard Medical School, 55 Fruit Street, Boston, Massachusetts 02114 USA; Precision Cardiology Laboratory, The Broad Institute, Cambridge, MA 02124 USA; Cardiovascular Developmental Biology Center, Department of Regenerative Medicine and Cell Biology, College of Medicine, Children’s Research Institute, Medical University of South Carolina, 171 Ashley Avenue, Charleston, SC 29425, USA; Inserm U1087; institut du thorax; University Hospital Nantes, France; CNRS, UMR 6291, Nantes, France; Université de Nantes, Nantes, France.; Department of Medicine, Division of Cardiology, University of California, San Francisco, 94143; Cardiac Ultrasound Laboratory, Cardiology Division, Massachusetts General Hospital, Harvard Medical School, 55 Fruit Street, Boston, Massachusetts 02114 USA; Center for Human Genetic Research, Massachusetts General Hospital and Department of Neurology, Harvard Medical School, 185 Cambridge St., Boston, MA 02114 USA.; Assistance Publique ‒ Hôpitaux de Paris, Department of Cardiology, Hôpital Européen Georges Pompidou, 75015, Paris, France; Assistance Publique ‒ Hôpitaux de Paris, Department of Genetics, Hôpital Européen Georges Pompidou, 75015, Paris, France

**Keywords:** heart valve disease, valve development, mitral valve prolapse, GWAS based enrichment analysis

## Abstract

Nonsyndromic Mitral valve prolapse (MVP) is a common degenerative valvular heart disease with severe health consequences, including arrhythmia, heart failure and sudden death. MVP is characterized by excess extracellular matrix secretion and cellular disorganization which leads to bulky valves that are unable to co-apt properly during ventricular systole. However, the triggering mechanisms of this process are mostly unknown. Using pathway enrichment tools applied to GWAS we show that genes at risk loci are involved in biological functions relevant to cell adhesion and migration during cardiac development and in response to shear stress. Through genetic, *in silico* and *in vivo* experiments we demonstrates the presence of several genes involved in gene regulation, including *GLIS1*, a transcription factor that regulates Hedgehog signaling. Our findings define genetic, molecular and cellular mechanisms underlying non-syndromic MVP and implicate disrupted endothelial to mesenchymal transition and cell migration as a potential common cause to this disease.

---

Mitral valve prolapse (MVP) is a common heart valve disease with important health consequences and an estimated prevalence of 2.4% in general populations^1,2^. It is defined as an abnormal mitral leaflet displacement into the left atrium during systole^3^ and is the most common and an increasing indication for surgical repair of mitral regurgitation (MR)^4,5^. MVP is a risk factor for heart failure and sudden death in community-based studies^1,4,6^. Prospective studies have showed that asymptomatic patients with low-risk presentation (e.g., moderate MR and ejection fraction ≥50%) can develop adverse MVP-related events, indicating wide heterogeneity in outcomes among individuals^4^.

The mature valve structure represents the culmination of embryonic programs working together to create cellular and extracellular matrix (ECM) environments that can withstand the biomechanical stresses of repetitive cardiac motion ^7^. These programs include early, growth factor-mediated endothelial to mesenchymal transition (EndoMT), followed by ECM organization and leaflet thinning. The mature postnatal leaflet structure is organized into stratified ECM layers of dense fibrous collagen in the ventricularis, a proteoglycan-rich central “spongiosa” and an elastrin-rich atrialis. Endothelial cells line the valve tissue whereas fibroblastic-like ECM-producing valve interstitial cells (VICs) cells make up the bulk of the valve^7^. In MVP, the normal layered structure is lost, with expansion of myxoid-like extracellular matrix (ECM) that includes fragmented collagen and elastin and overabundance of glycosaminoglycans. This renders the valve incompetent to withstand the normal physiological stresses of the beating heart and failure of normal leaflet apposition with prolapse into the left atrium. The process that leads to MVP and myxomatous degeneration is poorly understood. Previous reports have suggested that a potential mechanism is activation of quiescent VICs to myofibroblasts. These activated cells secrete matrix metalloproteinases that drive collagen and elastin fragmentation and release TGF-β that in turn promotes further cell proliferation and myofibroblast differentiation^7^. However, the triggering mechanisms of myxomatous degeneration are still to be identified.

Previous familial and population genetic studies have contributed to the current understanding of MVP biology. The identification of mutations in the filamin A gene (*FLNA)*^8^ followed by research in mice^9^ confirmed the role of this actin-binding protein during fetal valve development, mainly by providing stability to F-actin networks and linking them to cell membranes, which protect cells against mechanical stress^10^. More recently, we found that loss of function mutations in *DCHS1*, coding a protein from the cadherin superfamily involved in cell adhesion, cause familial MVP^11^. *DCHS1* deficiency in VICs altered migration and cellular patterning from patients and provoked loss of cell polarity during valve development in mice and zebrafish models^11^. Additional clues to the etiology of MVP came through genome-wide association studies (GWAS) where we identified six risk loci that are robustly associated with genetic susceptibility to MVP^12^. Genetic and biological evidence supported the role of two genes; tensin 1 (*TNS1*), a focal adhesion protein involved in cytoskeleton organization, and a transcription factor called LIM and cysteine-rich domain 1 (*LMCD1*)^12^. However, biological functions and potential mechanisms at the remaining GWAS loci remain unknown.

In this study, we hypothesize that several loci, including those not prioritized according to the stringent GWAS statistical threshold (P-value<5×10^−8^), could contain biologically relevant genes and mechanisms to MVP. We applied several computational-based analytical methods to the GWAS findings to 1) highlight enriched biological mechanisms for MVP loci, 2) characterize their expression pattern in tissues and 3) identify biologically pertinent genes for follow-up. Our study provides evidence for the Hedgehog signaling component, *GLIS1* to be involved in MVP susceptibility and provides supporting experimental validation of this gene in valve development and the degenerative process.

## Results

### Gene-set enrichment analyses for MVP loci

We first applied the SNP ratio test^13^ (SRT) method that uses simulated datasets to estimate the significance of a given pathway on the currently available GWAS data^12^. We analyzed ~2 million SNPs that mapped within genes and found 42 nominally enriched pathways (*P*_empirical_< 0.05) (Supplementary Table 1). Among the top 10 enriched pathways we found several pathways that are consistent with cardiac function, including viral myocarditis (*P*_empirical_<0.001), hypertrophic cardiomyopathy (*P*_empirical_<0.002), dilated cardiomyopathy (*P*_empirical_<0.002) and cardiac muscle contraction (*P*_empirical_<0.005). We also noted the enrichment in the phosphatidylinositol signaling system (*P*_empirical_<0.006), MAPK (*P*_empirical_<0.05) and WNT (*P*_empirical_<0.05) signaling that are important for focal adhesion and the gap junction functions (*P*_empirical_<0.05).

We next employed i-GSEA4GWAS, which highlighted 244 pathways as significantly enriched (FDR < 0.05) for MVP (Supplementary Table 2) and uses a larger source of pathways and gene sets compared to SRT (BioCarta, KEGG and Gene Ontology Biological processes). The most enriched pathway was BioCarta EDG-1 pathways with 26 out of 27 genes harboring associated variants to MVP. Globally, enriched pathways can be functionally classified into two main groups related to valve biology. The first and largest group included 28 GO terms related to ‘cytoskeleton’, ‘actin binding’, ‘focal adhesion’, ‘adhesion junction’ and ‘basolateral plasma membrane’, BioCarta gene set ‘Integrin pathway’ and KEGG pathway ‘cell adhesion molecules’. The second group was related to biological gene sets involved in the development of cardiovascular system and included 21 GO terms, notably ‘heart development’, ‘vasculature development’, ‘regulation of heart contraction’, ‘cardiac muscle contraction’, and ‘angiogenesis’. Interestingly, many enriched gene sets covered functions related to gene regulation, specifically, ‘transcription and regulation’ (60 related pathways), e.g. ‘transcription repressor activity’, ‘positive regulation of cell proliferation’, ‘positive regulation of transcription’. We note that this method also highlighted several consistent pathways with the enrichment obtained by the SRT method (e.g. dilated cardiomyopathy).

We also applied the integrative method *DEPICT* and used the recommended association cut-off (P_GWAS_-value <10^−5^) that identified 39 independent MVP loci harbouring 50 genes. We obtained 309 nominally enriched gene sets for MVP (P-value<0.05) and the Pearson distance matrix provided by DEPICT defined 36 clusters from these enriched gene sets (Supplementary Table 3). The second network is linked to developmental cardiovascular phenotypes which core node is cluster 3: vasculature development. This cluster contains mainly ‘abnormal blood vessel morphology’, ‘abnormal outflow tract development’, ‘failure of heart looping’, ‘angiogenesis’, ‘dorso-ventral axis formation’. Other clusters in the second network were cluster 5: ‘abnormal semilunar valve morphology’, cluster 9: ‘TGF-β receptor binding’ and ‘response to fluid shear stress’, and cluster 13: ‘abnormal fourth branchial arch artery morphology’ and ‘chromatin DNA binding’. Consistently, cluster 5, cluster 9 and cluster 13 were also significantly enriched for MVP genes and many gene sets included transcription factors as the most contributing genes (e.g *LMCD1*, *RUNX1*, *FOXL1*, *GLIS1*) (Supplementary Table 3).

### Enrichment for expression in tissues and cell types

Based on reconstituted gene-tissue matrix generated by DEPICT, we analyzed the expression profile from arrays-based human transcriptomic data and found that 21 tissues or cell types were enriched for the expression of genes at MVP risk loci. The most enriched physiological system is the cardiovascular system with five significantly enriched tissues. Connective tissue, epithelial and stem cells, specifically mesenchymal stem cells and muscle were the most enriched cell types and tissues for the expression of MVP genes (Figure 1b, Supplementary Table 4).

**Figure 1.**
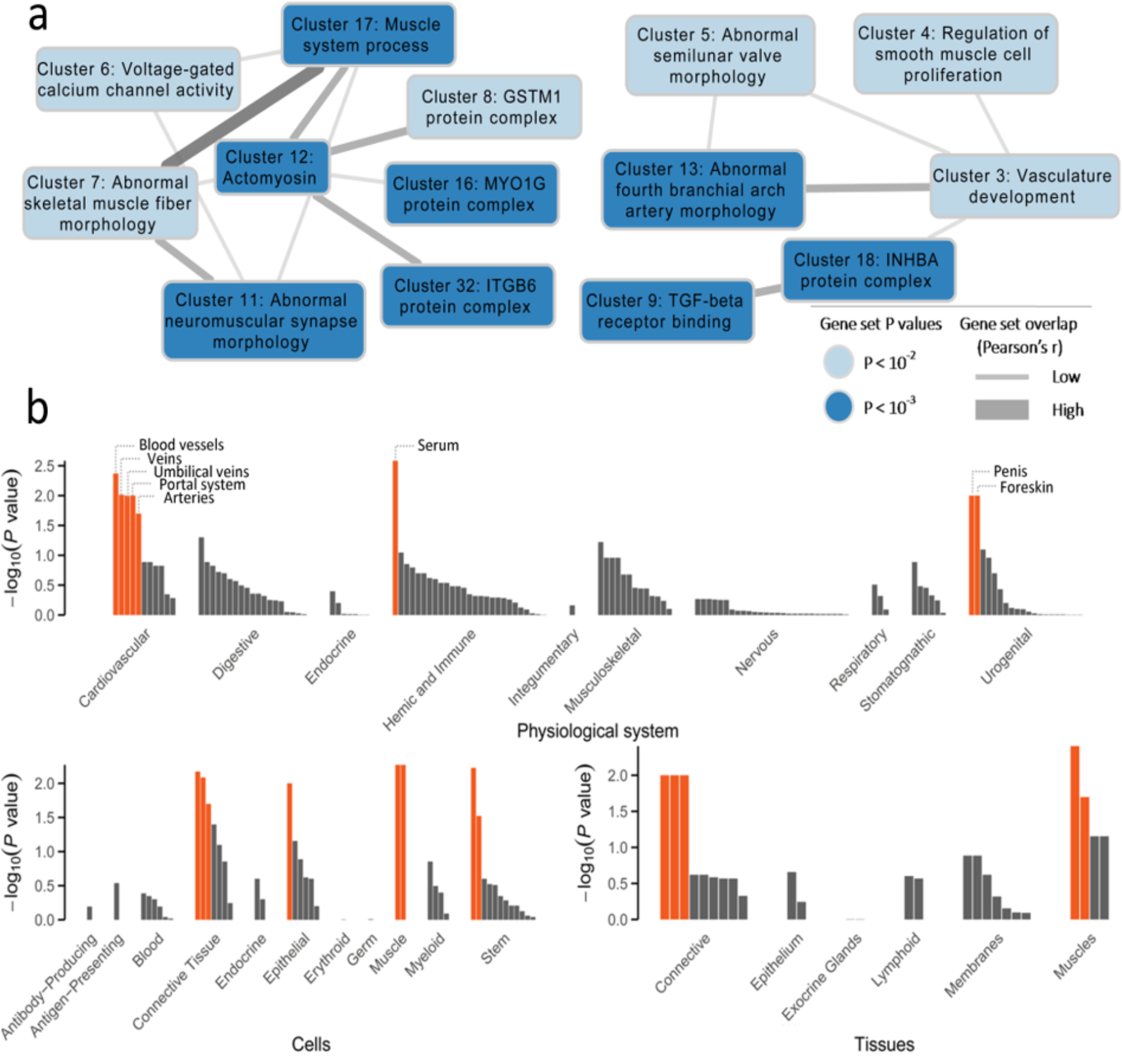
Networks of gene sets and tissue enrichment for the expression of genes at MVP associated loci. **(a)** Clustering networks formed from MVP enriched gene sets. The significance of the enrichment of gene sets are indicated (light blue: P-value < 10^−2^ and dark blue: P-value <10^−3^). The thicker the edge, the more similar functions exist between connected gene sets (all Pearson’s r > 0.3). (**b)** Tissue enriched for MVP genes. Enrichment is organised by physiological system, cells and tissues. Tissues/cells type with FDR < 0.20 are marked in orange.

### Prioritization of *GLIS1* as a novel risk locus and gene for MVP

#### Prioritization from contribution to enrichment analyses

Among the loci that we included in the pathway analyses, we provide a focus on a sub-GWAS significant signal located on chromosome 1 that deserved prioritization in the light of gene function candidacy and clues from the results obtained in the enrichment analyses. We found that i-GSWA4GWAS revealed *GLIS1* as the best-ranked gene in six significantly (FDR<0.05) and two suggestively (FDR<0.25, P-value<0.05) enriched gene sets, all related to regulation of transcription (Supplementary Table 5). Interestingly, Hh signaling pathway (from KEGG) was also associated with MVP (P-value=0.015, FDR=0.054, Supplementary Table 2). Although *GLIS1* is not reported as part of this pathway according to KEGG, several other genes from this pathway contain significantly associated variants with MVP (e.g *WTN5B*, *BMP6*, *HHIP*, *SHH*, *GLI2* and *GLI3* Supplementary Table 6). According to the DEPICT analysis, we found that *GLIS1* significantly contributes to the enrichment of several tissues and cell types namely the connective tissue (*GLIS1* Z-score=2.3), connective tissue cells (*GLIS1* Z-score=2.8) and mesenchymal stem cells (Z-score=2.3) (Supplementary Table 4). Of note, *GLIS1* is the most contributing MVP gene (Z-score=6.2) to suggestive enrichments (P-value=0.13, FDR>=0.20) of two cardiovascular system tissues (aortic valve and heart valves, Supplementary Table 4).

#### Genetic association at the GLIS1 locus

The association context at the *GLIS1* locus is presented in Figure 2a. The most associated SNP with MVP is rs1879734 and locates in the first intron (OR=1.30, P-value =7.1×10^−6^, risk allele: T, Freq: 0.28) (Supplementary Table 7). Follow-up of this SNP in four case control studies provided positive replication in the two largest studies, consistent direction of effect in all studies and no evidence for heterogeneity (OR_all_=1.23, P_effect_ =1.2×10^−7^, P_Heterogeneity_ = 0.41) (Supplementary Table 7).

**Figure 2.**
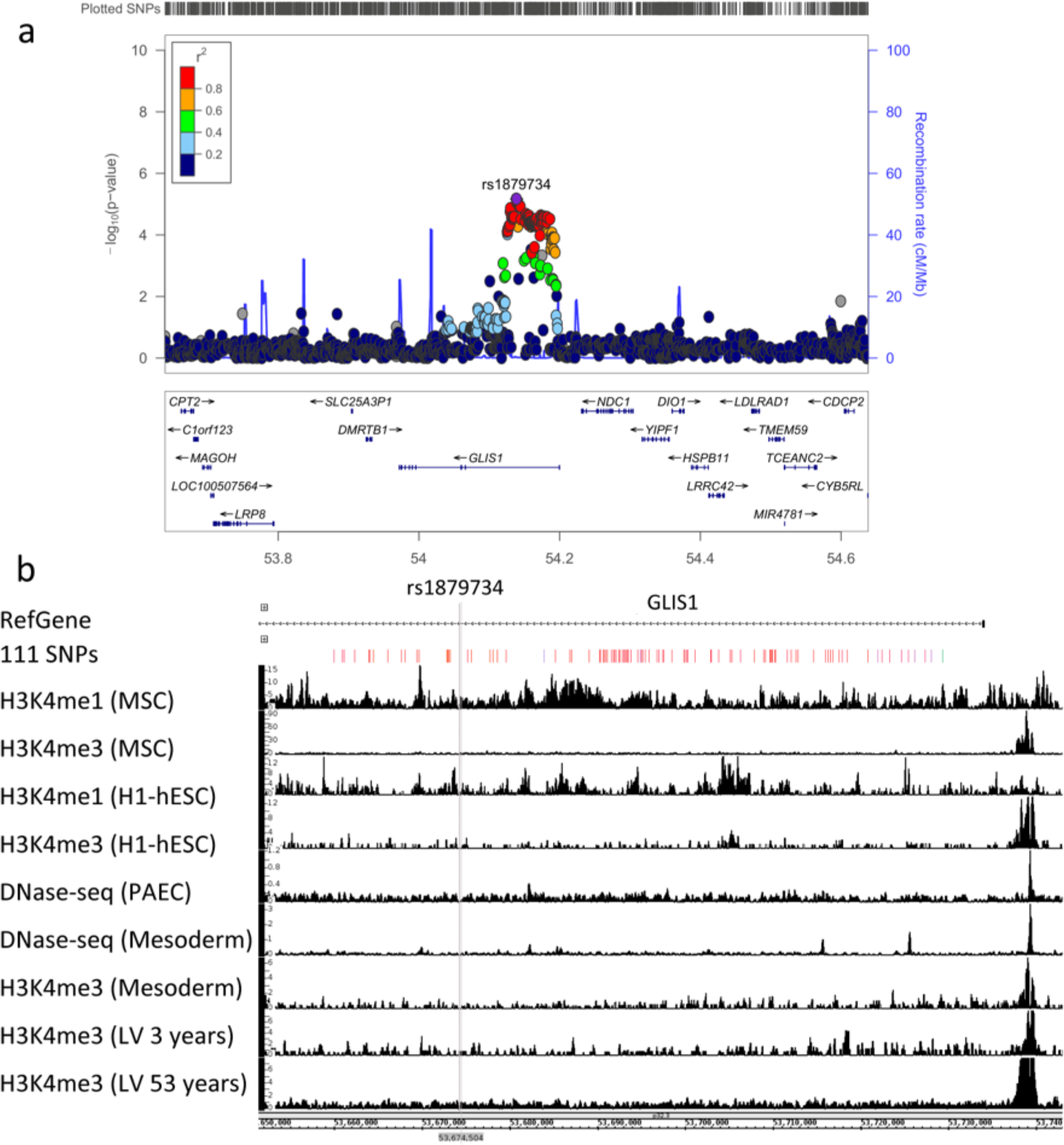
Genomic context and functional annotation of the association signal observed in the GWAS meta-analysis. **(a)** The regional association plot was generated using LocusZoom and displays surrounding genes (±500 Kb). The association signal is intronic to *GLIS1*. Round points represent SNPs in this region and purple point represent SNP rs1879734, the top associated SNP. **(b)** Visualization of histone marks and DNAse-seq density profiles in several tissues/cells based on ENCODE data. From top to bottom: reference gene; the selected 111 SNPs that is in high LD with rs1879734; H3K4me1 and H3K4me3 were from mesenchymal stem cell originated from H1-hESC; H3K4me1 and H3K4me3 from H1-hESC; DNase-seq from pulmonary artery endothelial cell; DNase-seq and H3K4me3 from cardiac mesoderm; H3K4me1 and H3K4me3 from heart left ventricle.

We performed functional annotation for 111 SNPs in moderate LD with rs18797434 (r^2^>0.5) that showed nominal association with MVP (P_GWAS_-value<0.05) and all mapped to introns1 and 2 in *GLIS1*(Supplementary Table 8). We found that 47 SNPs locate in DNase Hypersensitive region (DNASEV) in diverse normal tissues and 16 locate in transcription factor binding sites (TFBS). rs1879734 is located in a TFBS for endothelial transcription factor GATA2. Interestingly, we observed that rs12091931, which locates in DNASEV in non-pigmented ciliary epithelial cells, is in TFBSs for JUND, MAFF, MAFK, UBTF, MAZ, CTCF, ZBTB7A, TBP (Supplementary Table 8). Hereafter, the examination of the GTEx eQTL portal indicates that 58 SNPs, including the lead SNP rs1879734 were potential nominal eQTLs (P<0.05) for *GLIS1* in heart atrial appendage tissue (N=264). The most significant eQTL for *GLIS1* is rs2950241 (P-value=9×10^−3^), which is highly correlated to rs1879734 (r^2^=0.96, Hapmap CEU).

The examination of histone marks (Figue 2b) in mesenchymal stem cell originated from H1-hESC shows the presence of robust enhancer mark (H3K4me1) in the vicinity of rs1879734. In H1-hESC, strong signal of enhancer mark (H3K4me1) was presented at rs4927029, rs12097598, rs11206201, rs55786134, rs17109178. However, none of the 111 SNPs showed signals of histone marks in heart related tissue/cell types (pulmonary artery endothelial cell, cardiac mesoderm, heart left ventricle) (Figue 2b). *In silico* 4C experiment using four cell types including THP-1^14^, HUVEC^15^, H1-hESC^16^ and NHEK^15^ (Supplementary Figure 1) showed that the associated SNPs at this locus physically interact only with *GLIS1* regulatory sequences suggesting it is the potential target and causal gene at this locus.

### Expression during heart development in mouse

To study the pattern of expression of *GLIS1* during valve development, we performed immunohistochemistry (IHC) experiments of the mouse ortholog, *Glis1* during embryonic, foetal and adult time points (Figure 3). During embryonic development, Glis1 is expressed predominantly in nuclei of endothelial cells of the valves as well as the VICs. As the valves mature during foetal gestation, Glis1 is retained in a subset of endocardial and interstitial cells. By 6-months of age, Glis1 is much weaker in the valve leaflet with only scant cytoplasmic staining in endocardial cells. Weaker expression of Glis1 could be detected in the myocytes, epicardium and endocardium of the ventricular myocardium. These data show that Glis1 is embryonically expressed and that levels of this protein are rarely detected in the postnatal mouse, suggesting an important role for Glis1 in regulating valve morphogenesis during early development.

**Figure 3.**
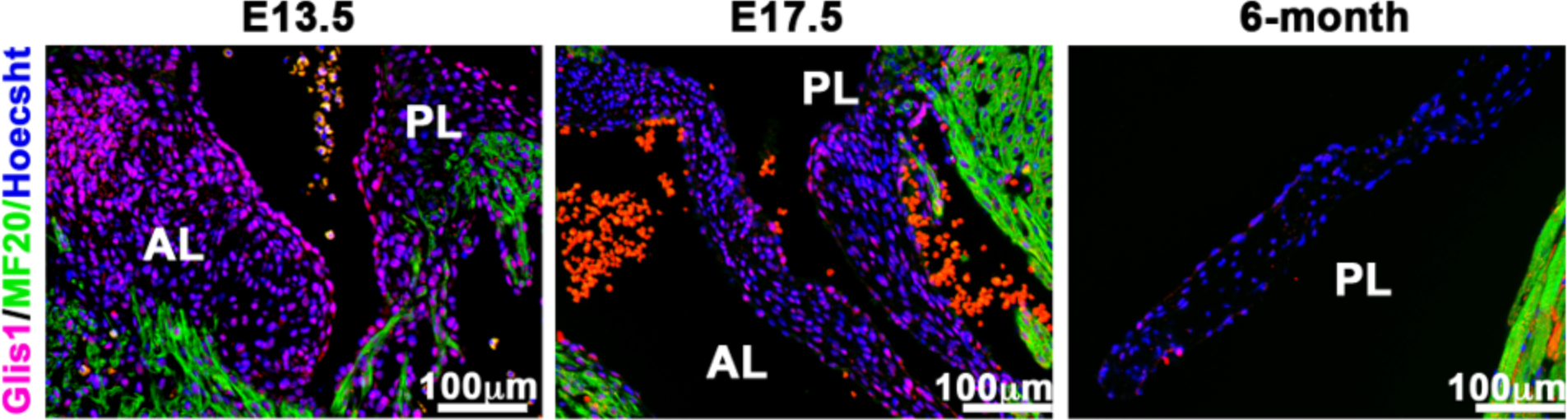
Glis1 expression in mouse developing and adult heart. Cardiac expression of Glis1 (red) was analysed at embryonic (E13.5), foetal (E17.5) and adult time points. Glis1 was detected during valve morphogenesis in mice (E.13 and E.17 stages), specifically during the completion of endothelial to mesenchymal transformation and valve sculpting and elongation and undetected in the adult valve (6 months). Glis1 is detected in nuclei from endothelial and valvular interstitial cells. Green tags are for MF20 marking sarcomeric myosin-myocytes, Blue is Hoescht coloration that indicates nuclei.

### Knockdown of *Glis-1* cause atrioventricular regurgitation in zebrafish

To analyse the potential effects of *GLIS-1* on valvular development and function, we chose to investigate its expression in the zebrafish. Due to genome duplication, zebrafish are predicted to have two orthologues of GLIS-1, *glis1a* and *glis1b*. We designed antisense morpholino oligonucleotides to target splice junctions in each with aims of rendering the transcript non-functional. Knockdown of *glis1a* was 65% efficient at 72 hours post fertilization, but had no discernible effect on atrioventricular valve function (Supplemental Figure 2). Knockdown of *glis1b* at 72 hours post fertilization was robust but slightly less efficient (37.3% and 36.5% reduction Figure 4a, b) and had minimal effects on the overt morphology of the developing embryo (Figure 4c). However, this knockdown resulted in a significant increase in the incidence of severe atrioventricular regurgitation when compared to controls. This increase was observed with two independently injected morpholinos with a combined fold increase of 1.6 (P=0.01) (Figure 4d).

**Figure 4.**
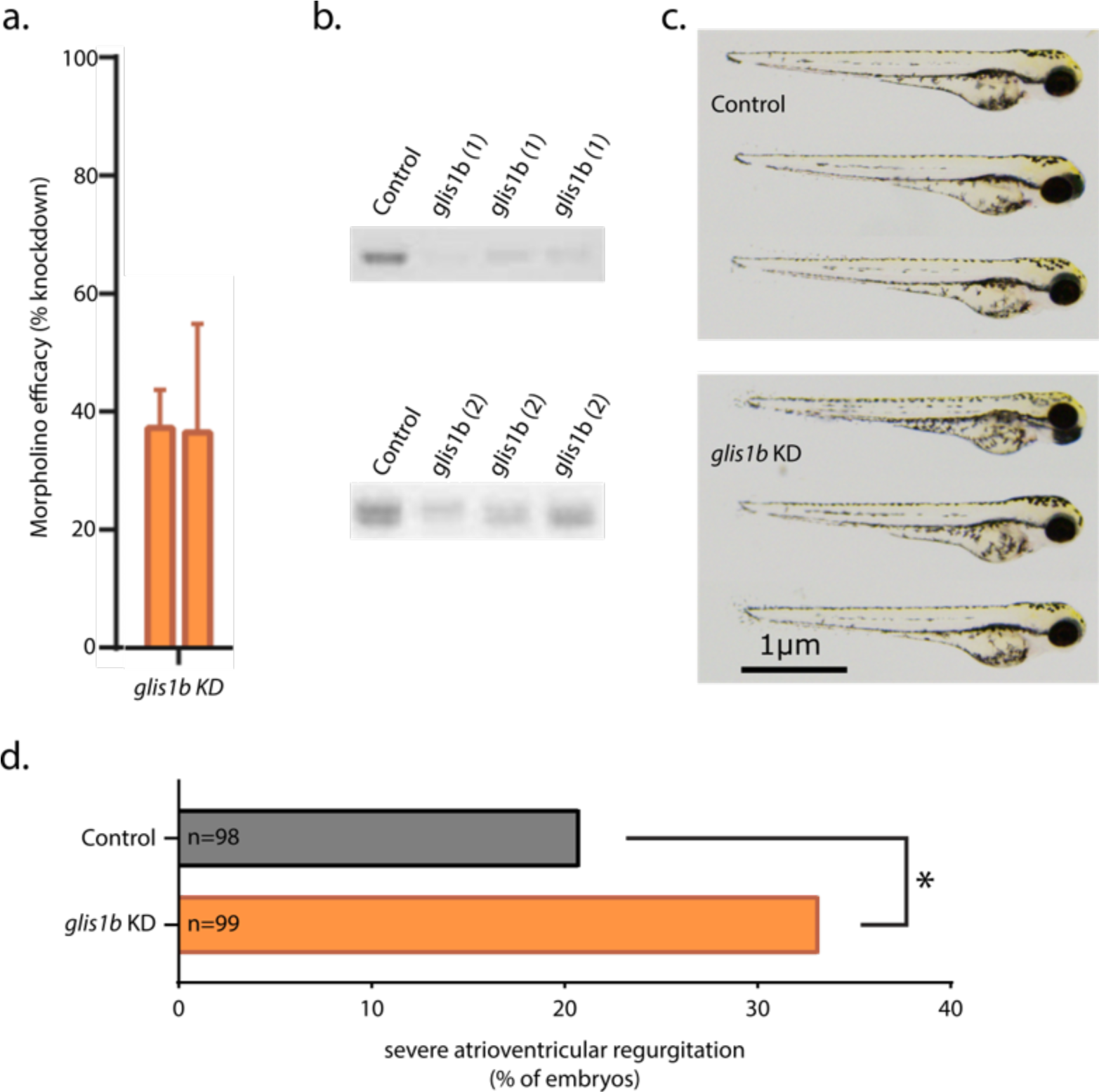
Assessment of cardiac regurgitation in zebrafish after morpholino knockdown of *glis1b.* **(a)** Morpholino-mediated knockdown efficacy. Efficacy for glis1b in embryonic zebrafish was measured by RT-PCR. **(b)** Representative gel images from analysis of morpholino efficacy. Control indicates samples amplified from control-injected embryos. All samples were obtained from 72-hpf embryos. **(c)**. Brightfield micrographs displaying gross morphology of 72-hpf embryos following glis1 knockdown. Scale bar represents 1 µm. **(d)** Fold change in observed atrioventricular regurgitation in 72-hpf zebrafish embryos after morpholino-mediated knockdown. All results are relative to clutchmate controls.

## Discussion

Our study supports that MVP genes are predominantly part of gene sets involved in cardiac development, the regulation of cell adherence and migration, and the regulation of cytoskeleton biology, focal adhesion and the interaction with the extracellular matrix. We also show that MVP genes are highly expressed in the cardiovascular system, connective and mesenchymal tissues. These genes are often regulatory genes expressed in the nucleus, as it is the case for *GLIS1*. Our findings suggest that this Hedgehog signalling related transcription factor is expressed in the nucleus of developing mitral valves in mouse and is required for atrioventricular development and function in zebrafish.

The application of three gene set enrichment tools to MVP GWAS data, using overall different computational methods and diverse databases provided overall consistent enrichments and an unprecedented resource to understand mitral valve biology. One of the most significantly enriched gene sets for MVP genes according to the i-GSEA4GWAS method is the endothelial differentiation gene (EDG-1) pathway where 26 out of 27 genes are associated with MVP. EDG-1 is a key signalling pathway for actin assembly and lamellipodia formation through the activation of integrins alpha V (ITGAV) and beta 3 (ITGB3) via RAS homolog gene family member A (RHOA). Edg1 knockout mice show embryonic haemorrhage leading to intrauterine death due to incomplete vascular maturation and defects in SPP-induced migration response^17^, which are important steps prior to valve sculpting at the embryonic stages. Cardiomyocyte specific knockout of EDG1 (alias S1PR1 for sphingosine 1-phosphate receptor-1) resulted in ventricular noncompaction and ventricular septal defects at 18.5 days post conception in mice, supporting a crucial role for this pathway in cardiac development^18^. Whether genes from EDG1 signalling are specifically important for valve development needs future examination. In support of this hypothesis is the reported interaction during lamellipodia formations of S1PR1 and Filamin A^19^, which gene is reported to be mutated in familial forms of MVP^20^. Many additional gene sets enriched for MVP loci are also involved in cardiac development or disease. These enrichments suggest critical roles for genetic variability near genes acting during early heart development that require important cytoskeleton remodelling and organization for cell migration during the endothelial-to-mesenchymal transition (EndoMT).

Our enrichment analyses also pointed to various pathways and gene sets whose link to valve disease is currently in its infancy. For instance, our analyses indicate an enrichment for gene sets in the immune system and phagosome pathways in MVP disease. The contention that the immune system may be involved in MVP is further supported by work from our group as well as others^9,21^. Additionally, the phagosome pathway has recently been reported as dysregulated in human myxomatous mitral valves compared with healthy valves^22^ and high activity of autophagosomes are present during cardiac development^23^. In addition, the knockdown of autophagy genes caused defects in cardiac looping and aberrant valve development as the results of ectopic expression of critical TF involved in heart development, mainly foxn4, tbx5 and tbx2^23^.

We report several top contributing genes at MVP loci to be transcription factors, especially to gene sets related to cardiovascular development. Consistently, we report enrichment for the protein localization to nuclear gene sets. Here, we followed-up specifically *GLIS1*, which was one of the top associated SNPs with MVP that we confirmed as a potential sub-GWAS risk locus for MVP. In addition to the expression in developing heart valves, *GLIS1* was the most significantly contributing gene to the enrichment of several gene sets related to the regulation of transcription. Little is known about GLIS1 function in connection with valve biology. We found that GLIS1 is expressed during developing valves in mouse valve endocardial and interstitial cells, suggesting for the first time its potential regulatory role during heart development. There is evidence for Glis1 to markedly increase the efficiency of generating induced pluripotent stem cells (iPSC) from mouse somatic cells in the presence of OCT3/4, SOX2 and KLF4^10^. This study demonstrated that GLIS1 directly interacts with KLF4 and induces the expression of Forkhead box genes, especially FOXA2 and several WNT genes to enhance mesenchymal to epithelial transition, a mechanism required for cell reprogramming^24^. The Hh signalling regulates both FOXA2 and WNT genes and its suggestive enrichment in our MVP GWAS involves for the first time in valve disease this important pathway for cell migration and morphogenesis organisation during heart development^25^. The Hh signalling induces EndoMT, through the up-regulation of NOTCH and TGF-ß signalling^26^ and is coordinated by primary cilia during cell migration and heart morphogenesis^25^. The contributing genes to the enrichment of the Hh signalling pathway included *BMP2* that harbours causative mutations for cardiac anomalies^27^ and *BMP4* that is required for outflow-tract septation^28^. Interestingly, a common variant in the also associated *WNT8A*, member of the WNT genes family is associated with atrial fibrillation^29^ and was previously shown to be regulated by GLIS1^10^.

This work presents however several limitations. One critical step when performing enrichment analyses in GWAS loci is the attribution of variants to genes. The pathway analyses of i-GWSEA4GWAS and DEPICT rely on SNPs mapped to genes using physical distance and LD block information. This have limited the number of genes analysed at risk loci and excluded more distant genes of interest. There is established evidence in favour of the functional role of distant long-range regulatory variants in predisposition to complex diseases^30^. In most cases, associated variants in GWAS loci are less likely to regulate the closest genes and be hundreds of kilobases away from culprit genes. This limitation explains the absence from the prioritization list generated by DEPICT of *TNS1*, a focal adhesion protein-coding gene that we have previously incriminated in MVP through genetic and functional investigation^12^.

In conclusion, our pathway investigation supports that genes near MVP associated loci are involved in biological functions relevant to cell adhesion and migration during cardiac development and in response to shear stress, and highlight the importance of regulatory mechanisms. Our study also provides genetic *in silico* and *in vivo* functional exploration of *GLIS1*, a transcription factor that regulate members of the Hh signalling and implicates this important biological mechanism for EndoMT and cell migration during heart development in valve myxomatous disease.

## Methods

### GWAS Study populations

The clinical characteristics of MVP patients, controls and GWAS methods have been previously described^12^. Briefly, we applied pathway-based methods to a meta-analysis of two GWAS conducted in 953 patients from MVP-France and 1566 controls and 489 from MVP-Nantes and 873 controls^12^. Replication was performed in four case-control studies (totalling 1,422 European MVP patients and 6,779 controls) as previously described^12^. Local ethics committees approved all studies, and all patients and controls provided written informed consent.

The input data was 6.6 million (M) genotyped or imputed SNPs (MAF > 0.01) and used either SNPTEST^31^ (imputed) or PLINK^32^ (directly genotyped) to perform the association test. Regional association plot at *GLIS1* locus was created using LocusZoom^33^.

### Pathway analyses

#### SNP ratio test (SRT)

SRT is a tool that calculates an empirical P-value by comparing the proportion of significant SNPs (P_GWAS_-value <0.05) in the original GWAS to randomized GWAS for the phenotype of cases and controls^13^. SRT tests enrichment using the KEGG pathways catalog and then allocates SNPs to genes using physical position and does not consider intergenic SNPs. Approximately 2 million GWAS SNPs were mapped to the reformed KEGG pathways and gene sets. We first generated 1000 simulated alternative phenotypes for individuals tested and ran the association analyses using PLINK^32^. The empirical P-value for a given pathway is defined by *P*_empirical_=*(s+1)/(N+1)*, where *s* is the number of times that a simulated ratio significant to non-significant was greater than or equal to the ratio obtained from the GWAS computed with real phenotypes, *N* is the total number of simulations (here *N*=1000) and enrichment was higher when P was small. We set significance to *P*-value<0.05 in simulated and real GWAS, and for the empirical P-value.

#### Gene set enrichment analyses using i-GSEA4GWAS

i-GSEA4GWAS v2 uses SNPs and their corresponding P-values from GWAS results as input^34,35^. All SNPs from the GWAS meta-analysis were used and were mapped to genes if they are exonic/intronic or to the closest genes if they are within 20 Kb upstream/downstream a gene. Of the 6.6M tested, 4.3M variants were mapped to 21,167 different genes and then genes were attributed to pathways and/or gene sets using BioCarta, KEGG, and GO terms from MSigDB v4.0. Only GO terms with experimental evidence (codes IDA IPI, IMP IGI, IEP), computational analysis evidence codes (ISS) and author evidence statement (TAS) were taken into account. In total, 936 gene sets contributed to calculate the significant proportion based enrichment score (SPES). We used the authors’ recommendation to consider pathways/gene sets with FDR<0.25 as suggestively associated with disease and FDR<0.05 as high confidence or statistically significant enrichment.

#### Integrative analyses using DEPICT

Data-driven expression prioritized integration for complex traits (DEPICT) (version rel194) is an integrative tool that utilizes diverse sources to predict reconstituted gene sets specific to each meta-analysis of GWAS^36^. DEPICT uses predefined gene sets including protein-protein interactions database, Mouse Genetics Initiative database, Reactome, KEGG and GO terms. It also uses a co-regulation data frame downloaded from the Gene Expression Omnibus (GEO) database including human and rodent expression microarrays (total arrays 77,840, including 37,427 generated in human tissues). This information was used to perform gene prioritization at loci, gene set enrichment analyses, and provide genes expression profiles in 209 tissues or cell types defined by Medical Subject Heading (MeSH). Input of DEPICT is a list of the most significantly associated SNPs (P_GWAS_-value<1×10^−5^) after LD pruning. Loci were mapped to genes using LD (r^2^>0.5). Genes where several SNPs map are counted once. To group overlapping gene sets, pairwise Pearson’s correlation coefficient of all enriched gene sets was calculated and the Affinity Propagation (AP) method^36,37^ was used to cluster gene sets that are highly correlated to automatically define independent clusters based on the Pearson distance matrix. We set significance for enrichment to P-value < 0.05, given that no gene sets reached an FDR<0.05.

### Variants Annotation

We used the UCSC genome browser tool Variant Annotation Integrator (VAI) to annotate SNPs at the *GLIS1* locus, and indicate if SNPs were located on DNase Hypersensitive region or transcription factor binding site generated by ENCODE^38^. Tissue specific expression quantitative trait loci (eQTL) annotation was extracted from GETx portal (https://www.gtexportal.org/home/, Release V7, dbGaP Accession phs000424.v7.p2). Annotation of the 111 selected SNPs by Human ChIP-Seq (markers: H3k4ME1 and H3k4ME3) and DNase-Seq were downloaded from ENCODE (https://www.encodeproject.org/) and visualized by Integrated Genome Browser (IGB)^39^. In the absence of valvular cells in this database, selected tissues/cells used in this analyses are mesenchymal stem cell originated from H1-hESC^40^, H1-hESC^40^, pulmonary artery endothelial cell^41^, cardiac mesoderm^41^ and heart left ventricle^40^. Accession numbers of those files are (respectively): ENCFF152YQG (H3K4me1), ENCFF712CJP (H3K4me3), ENCFF623ZAW (H3K4me3), ENCFF593OAZ (H3K4me1), ENCFF719ZEX (DNase-seq), ENCFF591TLE (H3K4me3), ENCFF213FJN (DNase-seq), ENCFF094USN (DNase-seq), ENCFF254JZR (DNase-seq). The annotation of the *GLIS1* region used the Hi-C data from four cell types including THP-1^14^, HUVEC^15^, H1-hESC^16^ and NHEK^15^ from ENCODE, all visualized by IGB.

### Protein detection in mouse embryos and adult hearts

In all IHC experiments, 5-min antigen retrieval was performed with VectaStain in a pressure cooker (Cuisinart). The antibodies used for IHC were: Glis1 (Novus, NB100-41087) and myosin heavy chain (Developmental Hybridoma Banks, MF20). Primary antibodies were used at a 1:100 dilution; Hoechst 33342 (nuclear stain) was used at a 1:10,000 dilution. Appropriate secondary antibodies were used for detection. Three time points were used in IHC: (i) completion of the endothelial-to-mesenchymal transition (EndoMT; embryonic day (E) 13.5), (ii) valve sculpting and elongation (E17.5) and (iii) achievement of the mature adult form (at 9 months).

### Zebrafish experiments

Zebrafish experiments were performed in accordance with approved Institutional Animal Care and Use Committee (IACUC) protocols. Zebrafish of the Tu-AB strain were reared according to standard techniques. Morpholinos were designed against the two zebrafish orthologues of GLIS1: *glis1a* (5’-GAGAATGGTCGTACATACCGTGTCC) and *glis1b* (1: 5’-AAGTGCACTGAGGTCTCACCCTGTG 2: 5’-TTAAGGTCAGGTACTCACAGTGTCC). Empirically established effective doses of antisense morpholino oligonucleotides were injected at the single-cell stage, and morpholino-injected animals were compared to controls injected with a non-targeting morpholino. Analysis of atrioventricular regurgitation was performed at 72 hours post fertilization as described^12^.

## Acknowledgments

This study was supported by a PhD scholarship from the Chinese Scientific Council to MY, and partly funded by French Agency of Research (ANR-16-CE17-0015-02). The work at MUSC was performed in a facility constructed with support from the National Institutes of Health, Grant Number C06 RR018823 from the Extramural Research Facilities Program of the National Center for Research Resources. Other funding sources: National Heart Lung and Blood Institute: HL131546 (RAN), COBRE GM103342 (RAN), GM103444 (RAN), HL127692 (DJM, SAS, RAN, RAL), American Heart Association: 17CSA33590067 (RAN) and HL140187 (NRT). The recruitment of the MVP France cohort was supported by the French Society of Cardiology. We acknowledge the contribution of the Leducq Foundation, Paris for supporting a transatlantic consortium investigating the physiopathology of mitral valve disease, for which this genome-wide association study was a major project (coordinators: R.A.L. and A.A.H.). We acknowledge investigatrs who contributed access to validation in cohorts: Leticia Fernandez-Friera, Jorge Solis from Centro Nacional de Investigaciones Cardiovasculares (CNIC), Yohan Bossé and Philippe Pibarot from *Institut universitaire de cardiologie et de pneumologie de Québec-Université Laval*, Quebec, Canada, Ramachandran S. Vasan, Ming-Huei Chen and Emilia J. Benjamin from the Framingham Heart Study, USA, Thierry Le Tourneau, Richard Redon, Hervé Le Marec and Vincent Probst from Institut du Thorax, Nantes, France and Ronen Durst, Hassadah Hebrew University Medical Center, Jerusalem, IL.

## Author contributions

Recruitment of patients: A.A.H, R.A.L, S.A.S, F.N.D, X.J.

Genotyping: X.J, J.-J.S, C.D, S.A.S.

Data analysis: M.Y, A.G, S.K

Animal experiments: D.J.M., R.A.N., N.T., P.T.E., K.T.

Manuscript writing: N.B.-N., M.Y, R.A.N., N.T., R.A.L.

Manuscript approval: all authors.

---

**Supplementary Figure 1.**
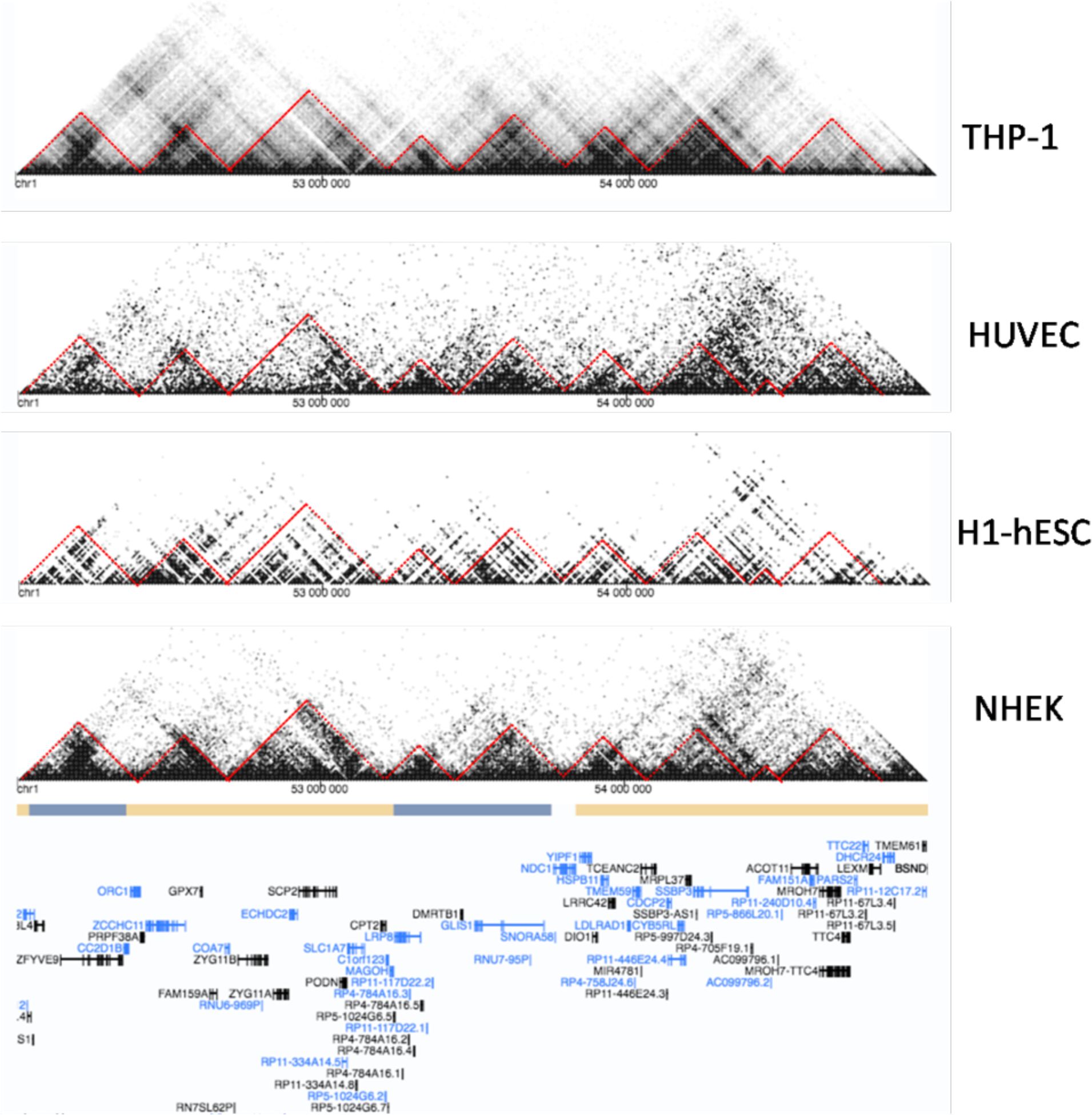
Hi-C interaction of GLIS1 in several cell types. We selected Hi-C data from four cell types includes THP-1, HUVEC, H1-hESC, NHEK from ENCODE, gene interaction visualized by Integrated Genome Browser (IGB) was shown. By visualization of the topologically associating domains (TADs), the interaction in the GLIS1 region was found to be mainly exists inside of GLIS1.

**Supplementary Figure 2.**
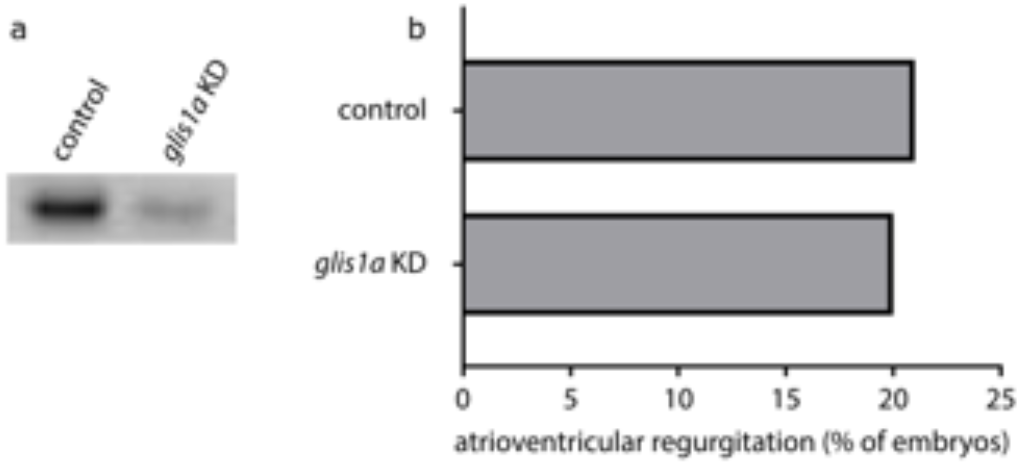
Assessment of cardiac regurgitation in zebrafish after morpholino knockdown of *glis1a*. **(a)** Representative gel images from analysis of morpholino efficacy. Control indicates samples amplified from control-injected embryos. All samples were obtained from 72-hpf embryos. (**b**) Fold change in observed atrioventricular regurgitation in 72-hpf zebrafish embryos after morpholino-mediated knockdown.

**Supplementary Table 1.** Enriched pathways for nominally associated loci in MVP GWAS according to the SNP ratio test (SRT) method. Enriched KEGG pathways are those for which the 1000 simulated GWAS with permuted case control status provided smaller counts of ratios of significant to non significant SNPs, when compared to GWAS with real phenotyes. We provide the count of ratios and empirical p-values (<0.05) for pathways. E.g Compared to the real GWAS, SNPs in genes in the phagosome pathway presented more significant associations with MVP only in 1 out of 1000 simulated GWAS. On the other hand, SNPs in genes from the Wnt signaling pathway presented more significant associations with MVP 50 simulated GWAS.

| Pathway ID in KEGG | Descriptive of the pathway | Count of ratios of significant to non-significant SNPs higher in simulated data | Empirical P value |
| --- | --- | --- | --- |
| hsa04145 | Phagosome | 1 | 0.001 |
| hsa05416 | **Viral myocarditis** | 1 | 0.001 |
| hsa05410 | **Hypertrophic cardiomyopathy (HCM)** | 2 | 0.002 |
| hsa05414 | **Dilated cardiomyopathy** | 2 | 0.002 |
| hsa00190 | Oxidative phosphorylation | 3 | 0.003 |
| hsa05340 | Primary immunodeficiency | 3 | 0.003 |
| hsa00010 | Glycolysis / Gluconeogenesis | 4 | 0.004 |
| hsa04612 | Antigen processing and presentation | 4 | 0.004 |
| hsa00051 | Fructose and mannose metabolism | 5 | 0.005 |
| hsa04260 | **Cardiac muscle contraction** | 5 | 0.005 |
| hsa04672 | Intestinal immune network for IgA production | 5 | 0.005 |
| hsa05140 | Leishmaniasis | 5 | 0.005 |
| hsa05310 | Asthma | 5 | 0.005 |
| hsa05330 | Allograft rejection | 5 | 0.005 |
| hsa05332 | Graft-versus-host disease | 5 | 0.005 |
| hsa01100 | Metabolic pathways | 6 | 0.006 |
| hsa04070 | **Phosphatidylinositol signaling system** | 6 | 0.006 |
| hsa00562 | Inositol phosphate metabolism | 7 | 0.007 |
| hsa05320 | Autoimmune thyroid disease | 7 | 0.007 |
| hsa04662 | B cell receptor signaling pathway | 8 | 0.008 |
| hsa04940 | Type I diabetes mellitus | 8 | 0.008 |
| hsa05210 | Colorectal cancer | 10 | 0.010 |
| hsa05144 | Malaria | 11 | 0.011 |
| hsa05322 | Systemic lupus erythematosus | 11 | 0.011 |
| hsa04660 | T cell receptor signaling pathway | 12 | 0.012 |
| hsa00740 | Riboflavin metabolism | 15 | 0.015 |
| hsa00750 | Vitamin B6 metabolism | 15 | 0.015 |
| hsa04370 | VEGF signaling pathway | 16 | 0.016 |
| hsa04930 | Type II diabetes mellitus | 18 | 0.018 |
| hsa04962 | Vasopressin-regulated water reabsorption | 18 | 0.018 |
| hsa00524 | Butirosin and neomycin biosynthesis | 25 | 0.025 |
| hsa04010 | **MAPK signaling pathway** | 30 | 0.030 |
| hsa00920 | Sulfur metabolism | 31 | 0.031 |
| hsa00603 | Glycosphingolipid biosynthesis - globo series | 36 | 0.036 |
| hsa04270 | Vascular smooth muscle contraction | 36 | 0.036 |
| hsa00350 | Tyrosine metabolism | 37 | 0.037 |
| hsa04540 | **Gap junction** | 41 | 0.041 |
| hsa04650 | Natural killer cell mediated cytotoxicity | 45 | 0.045 |
| hsa05215 | Prostate cancer | 45 | 0.045 |
| hsa00970 | Aminoacyl-tRNA biosynthesis | 48 | 0.048 |
| hsa04966 | Collecting duct acid secretion | 48 | 0.048 |
| hsa04310 | **Wnt signaling pathway** | 50 | 0.050 |

**Supplementary Table 2.**
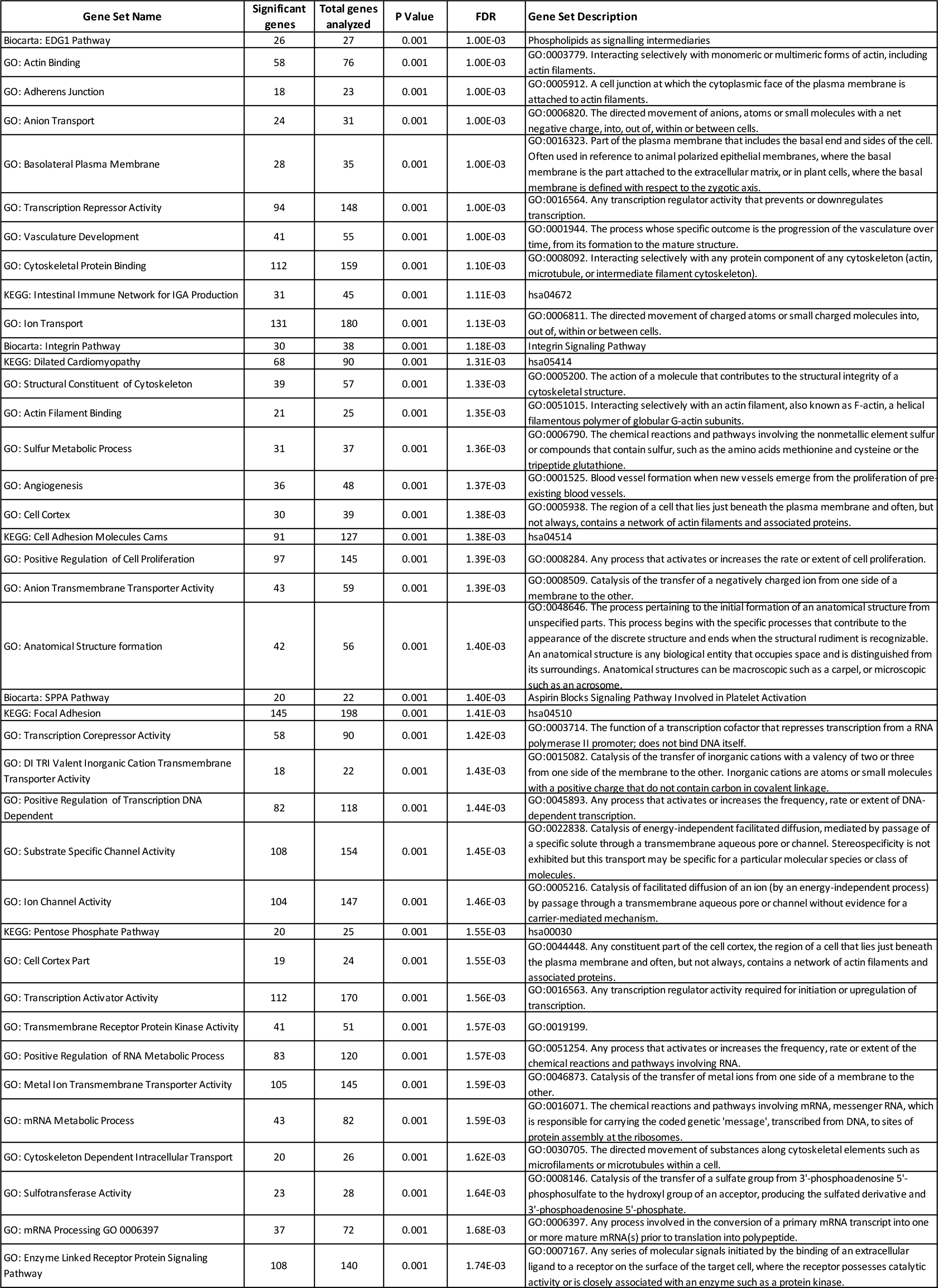
Enriched gene sets identified by i-GSEA4GWAS. We display all 244 confidently enriched pathways /gene sets (FDR < 0.05). We also added the Hedgehog Signaling Pathway result. The number of significantly contributing genes and the total number of genes analyzed are indicated. A short gene set description was provided when available, in addition to the catalog IDs (GO and KEGG). FDR: false discovery rate.

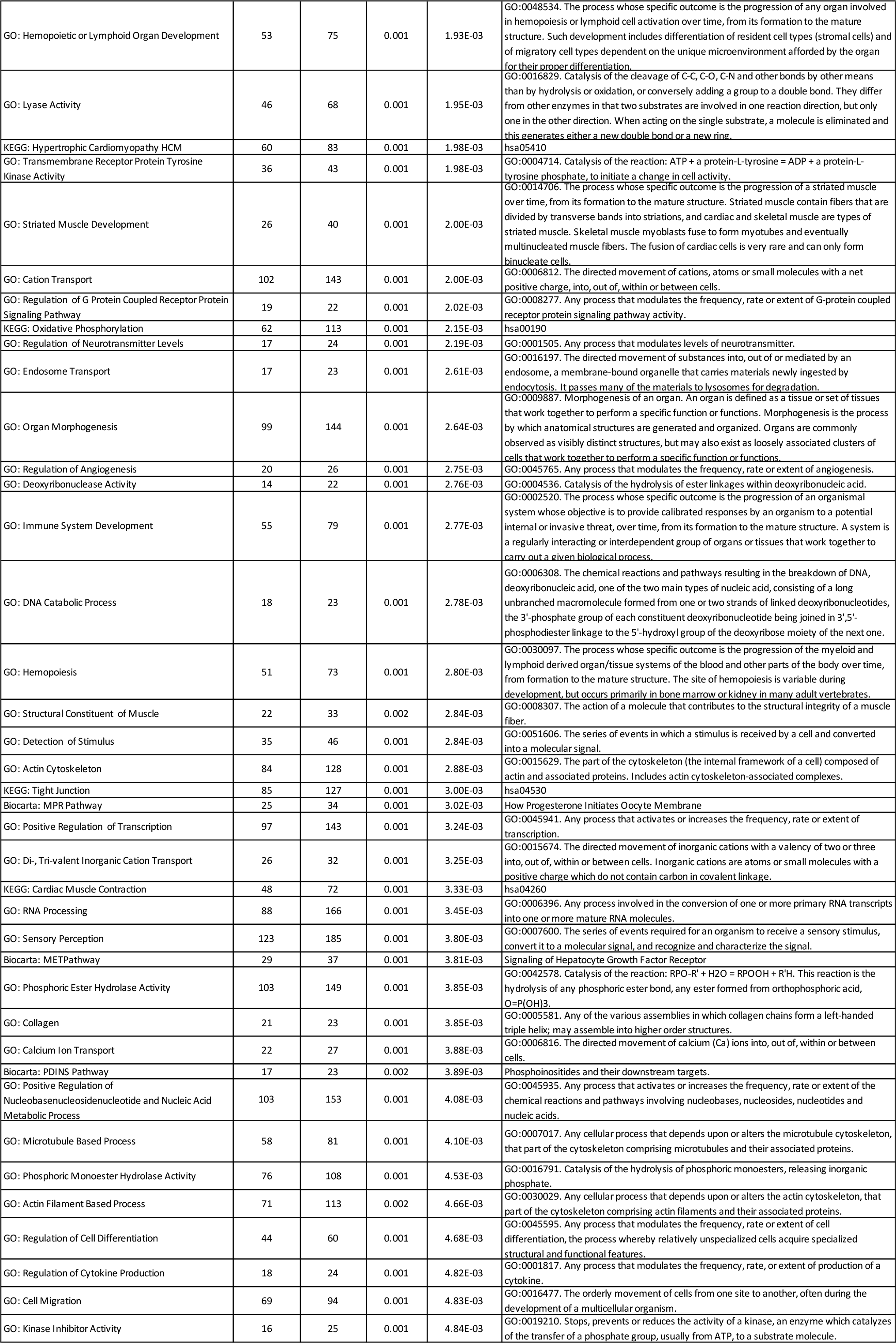

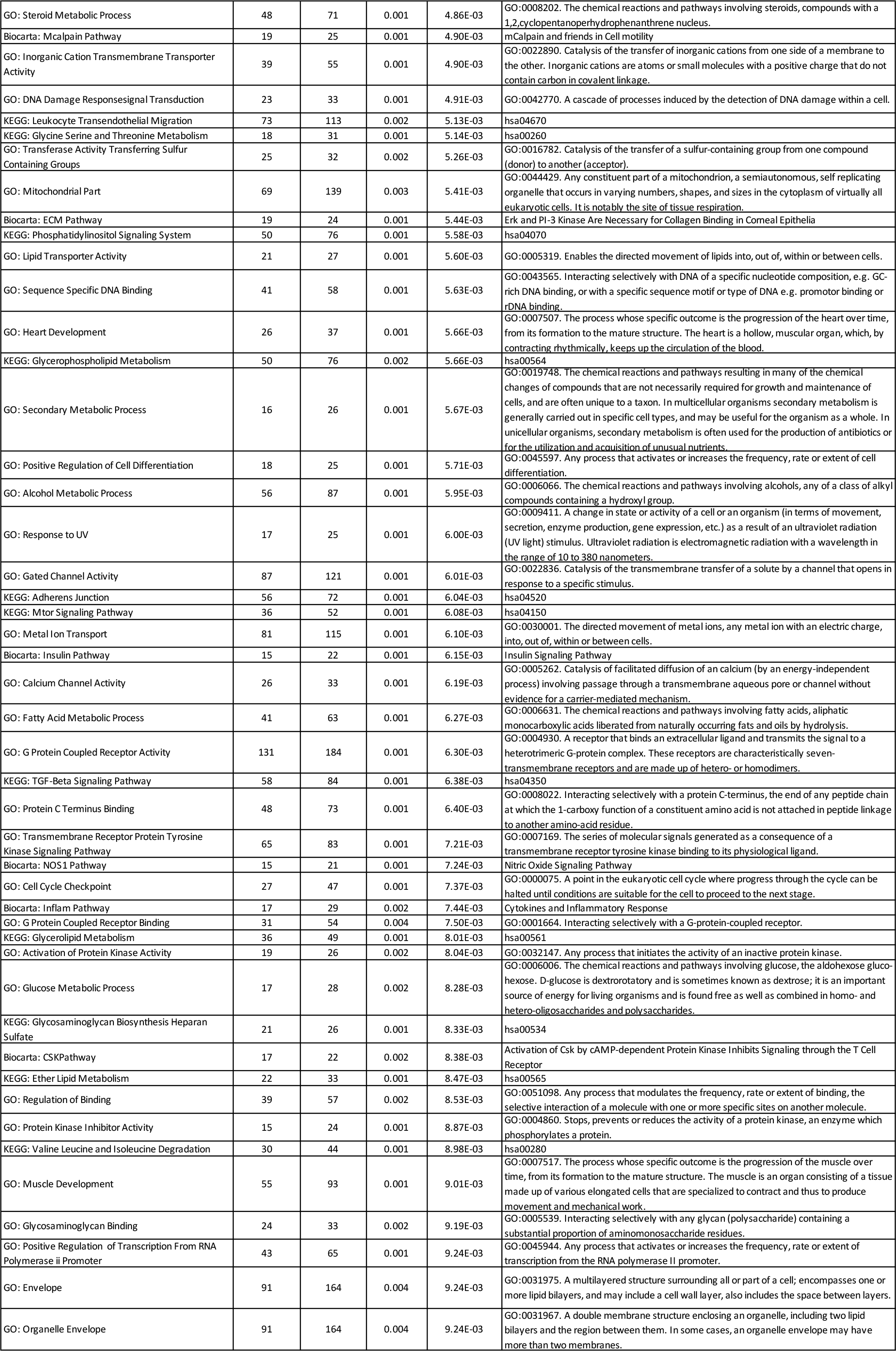

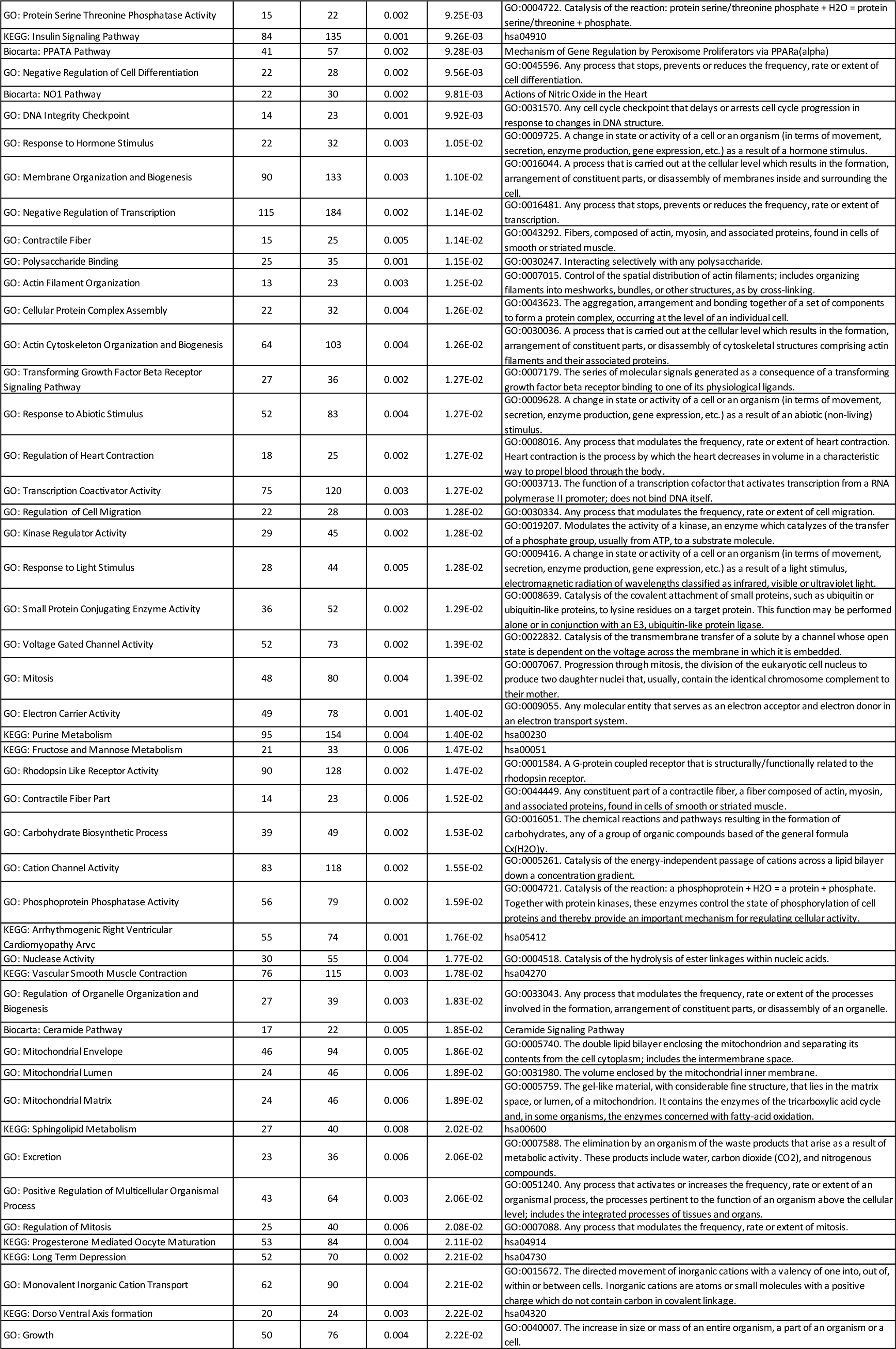

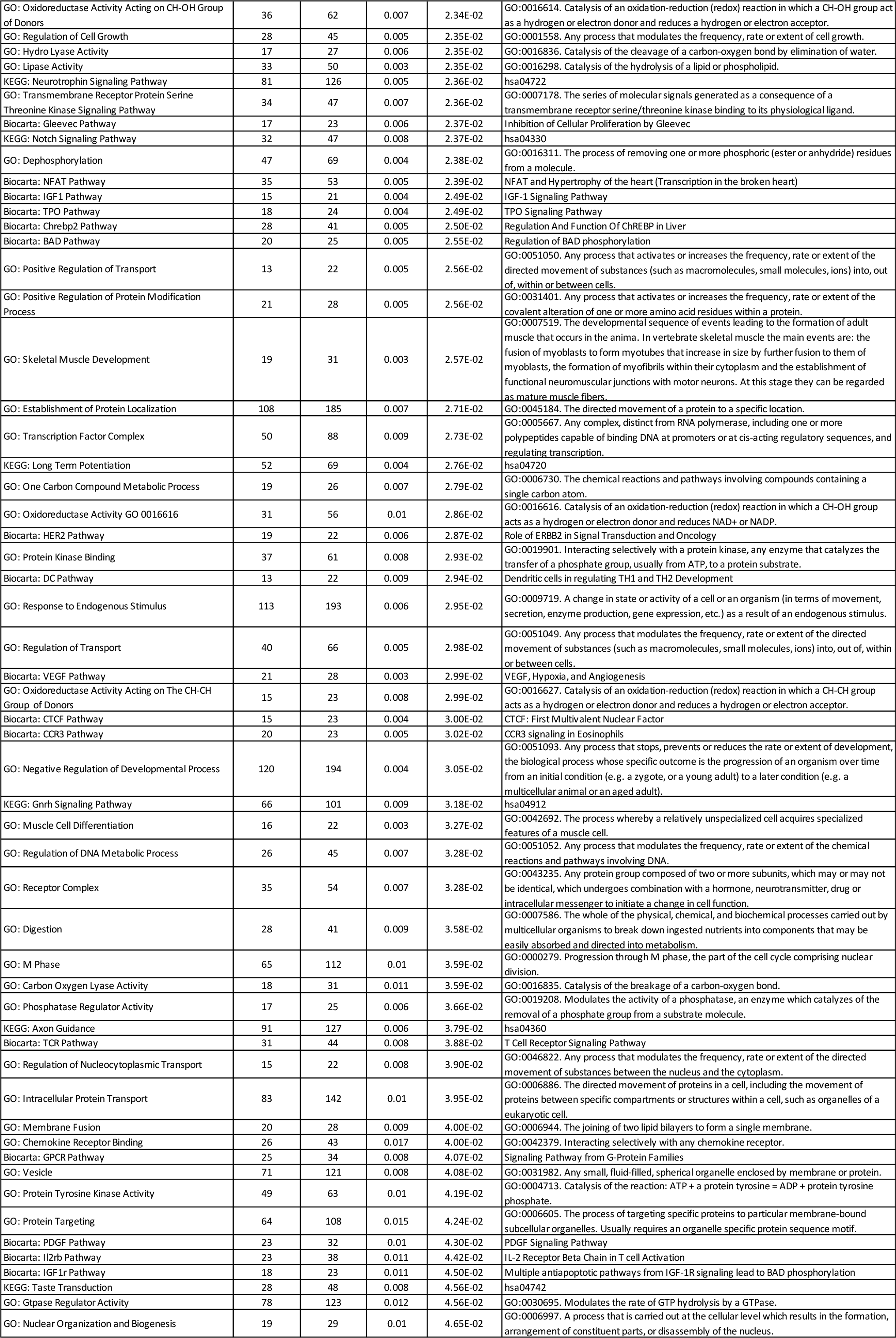

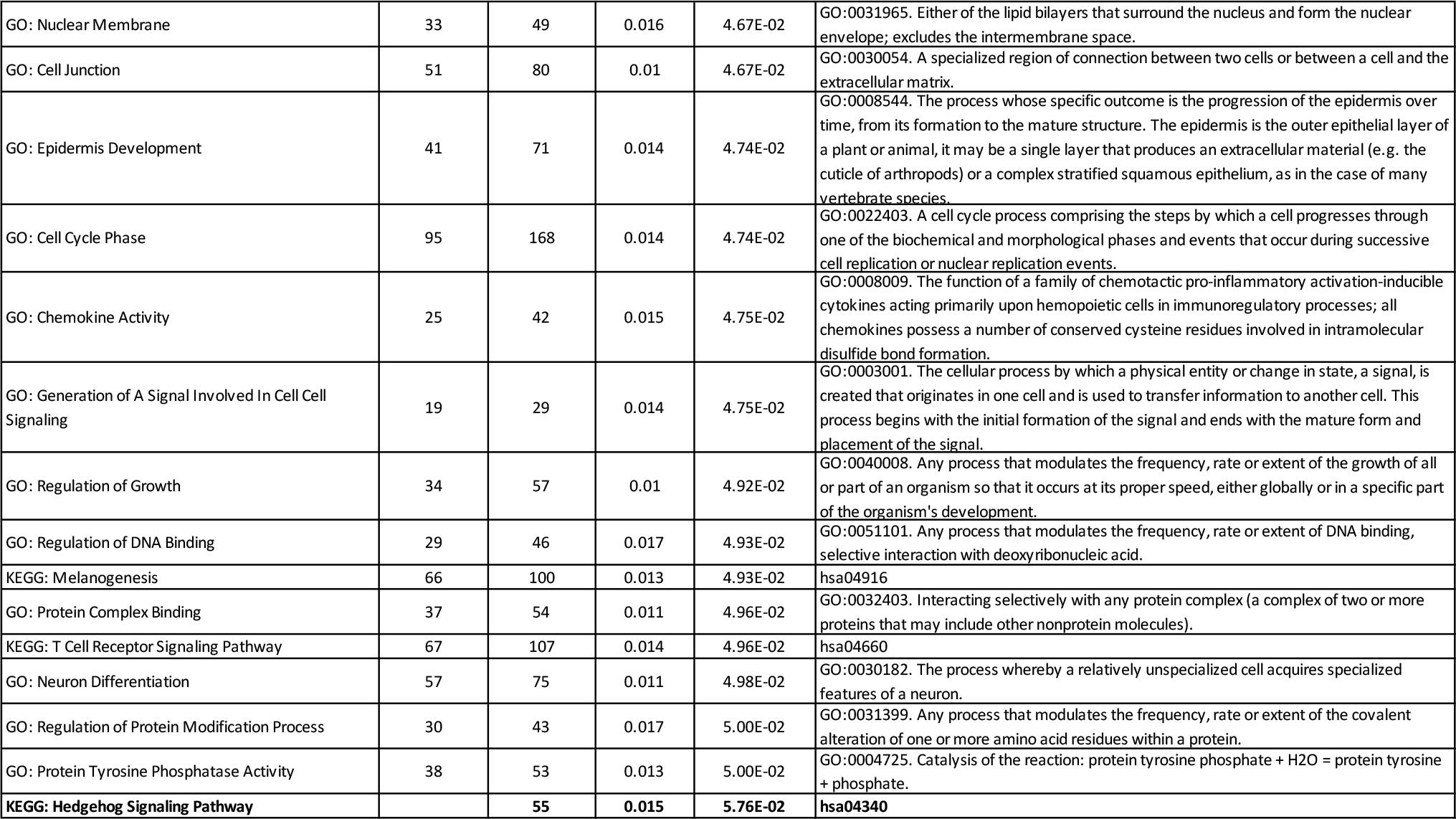

**Supplementary Table 3:**
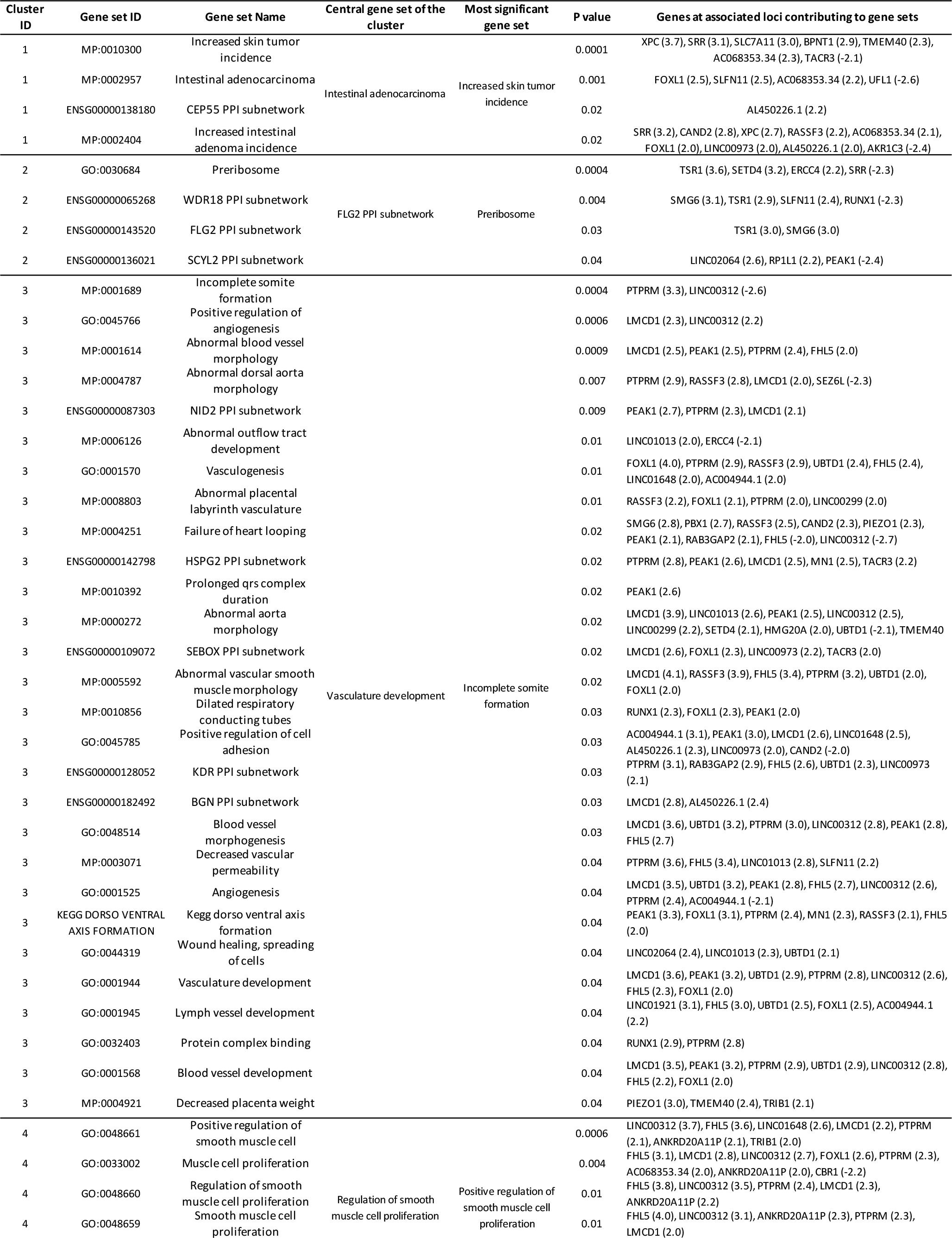
Enriched reconstituted gene sets identified by DEPICT. DEPICT was used to assess whether 50 genes located in associated regions (GWAS P-value < 1 × 10^−5^) were enriched for any of the reconstituted gene sets defined by DEPICT. We display 36 reconstituted clusters with at least one gene set that reaches suggestive enrichment (P-value < 0.01). For each cluster we provide all gene sets that contribute to this cluster and were nominally enriched (P-value<0.05). Associated regions were constructed by mapping genes to a given region if the genes resided within, or overlapped with linkage disequilibrium (r2 > 0.5) flanks of a given lead SNP. Ex: Cluster ID and cluster center was defined by Affinity Propagation method which was used to cluster gene sets that were highly correlated and automatically defines independent clusters based on calculated Pearson’s correlation matrix (See methods for dertails). The central gene set of a given cluster is the most significant gene set in that cluster (bold). We provide the list of genes near associated loci allocated to each gene set and their respective likelihood to be part of each gene set. We only provide genes with significant likelihood |z-score| > 1.96 corresponding to P-value <0.05. NA: not available, is indicated for gene sets for which no gene presented a significant z-score.

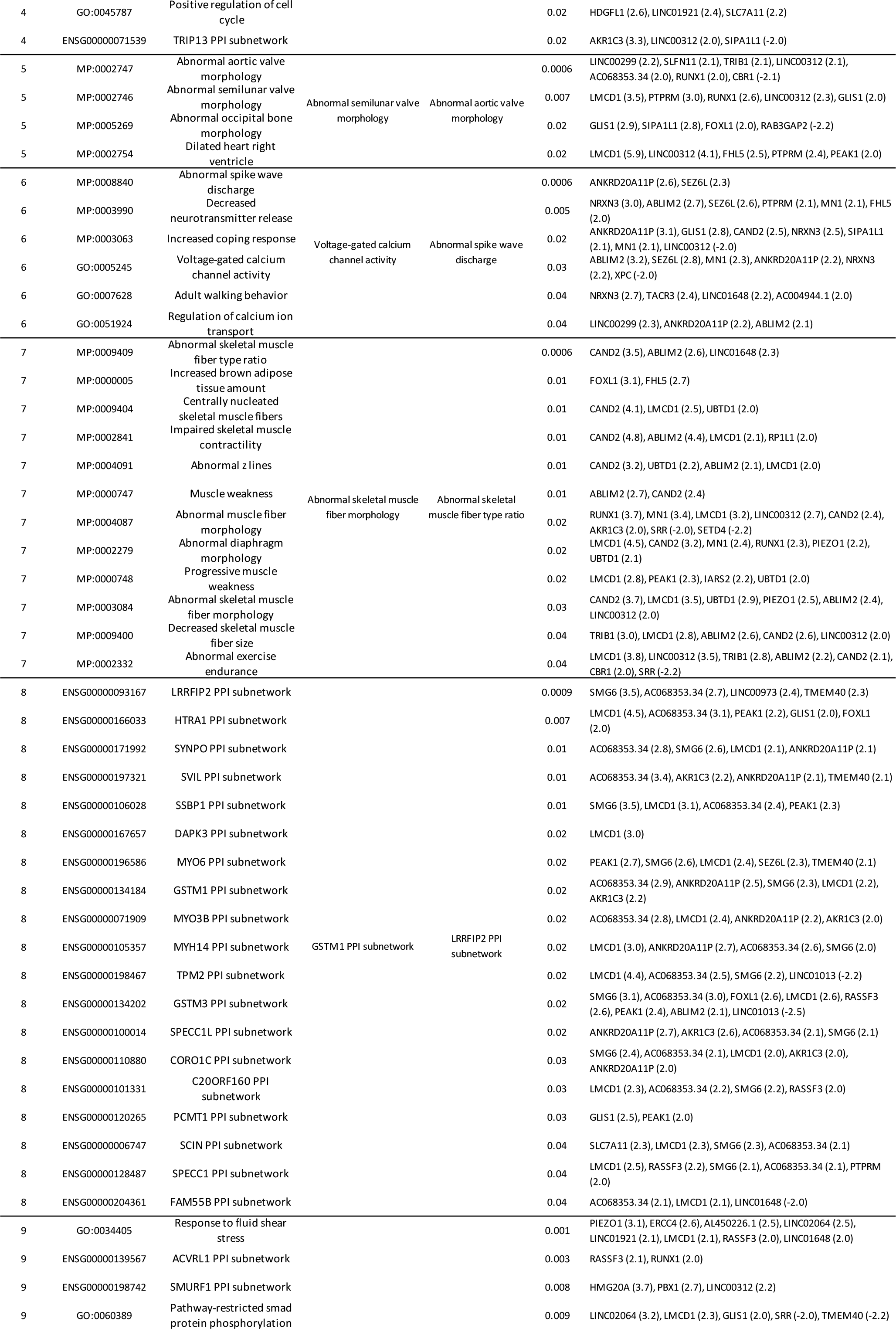

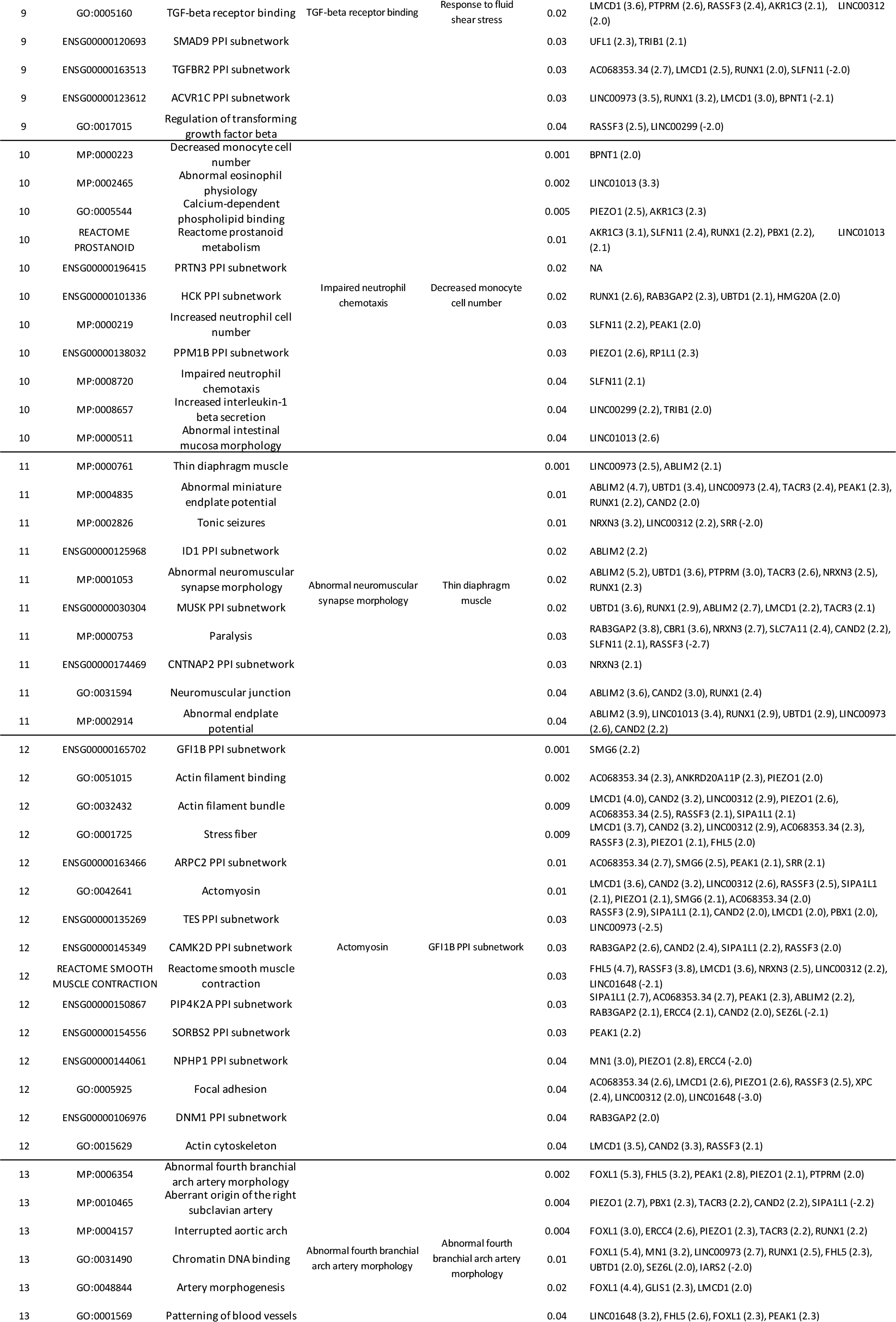

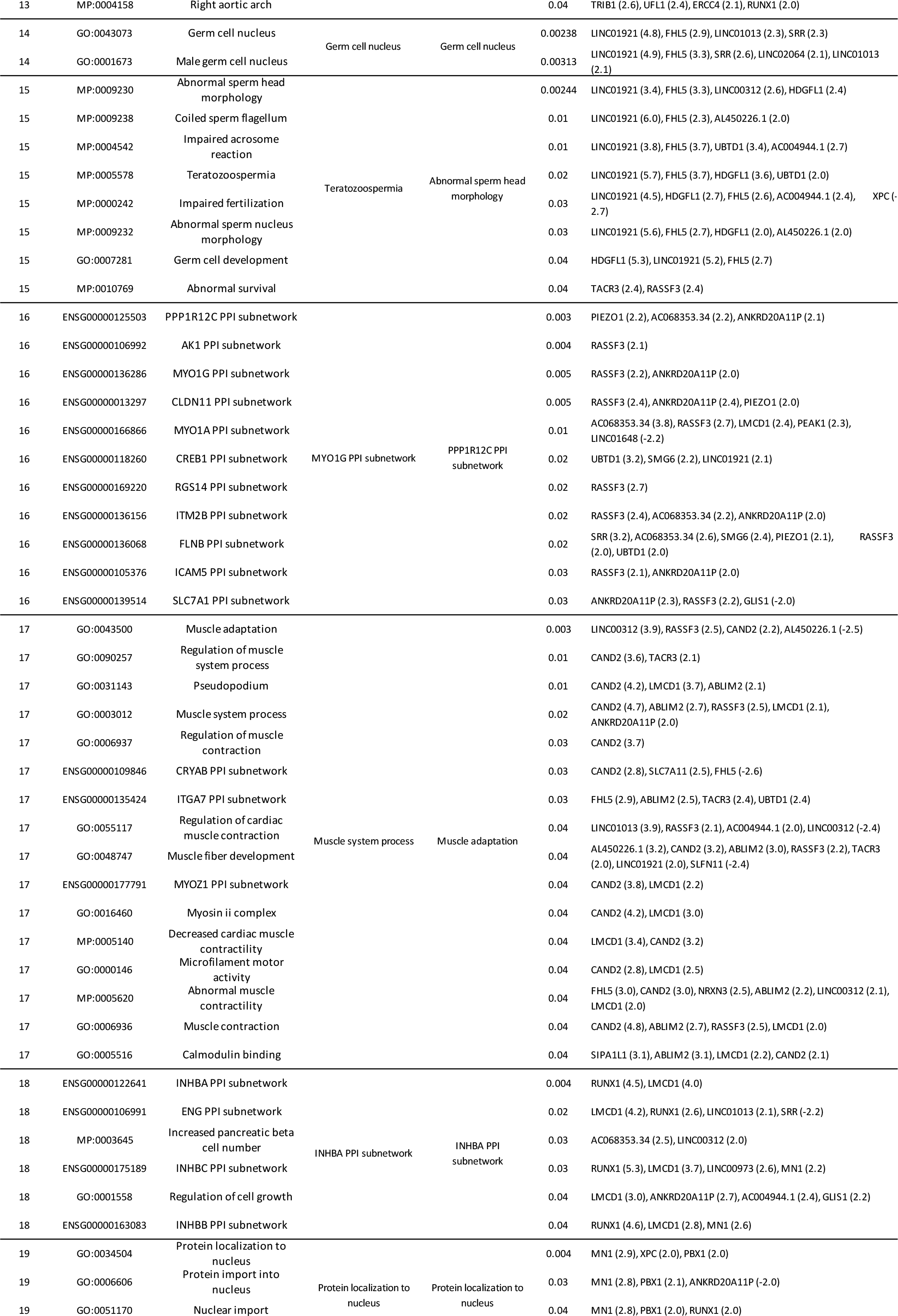

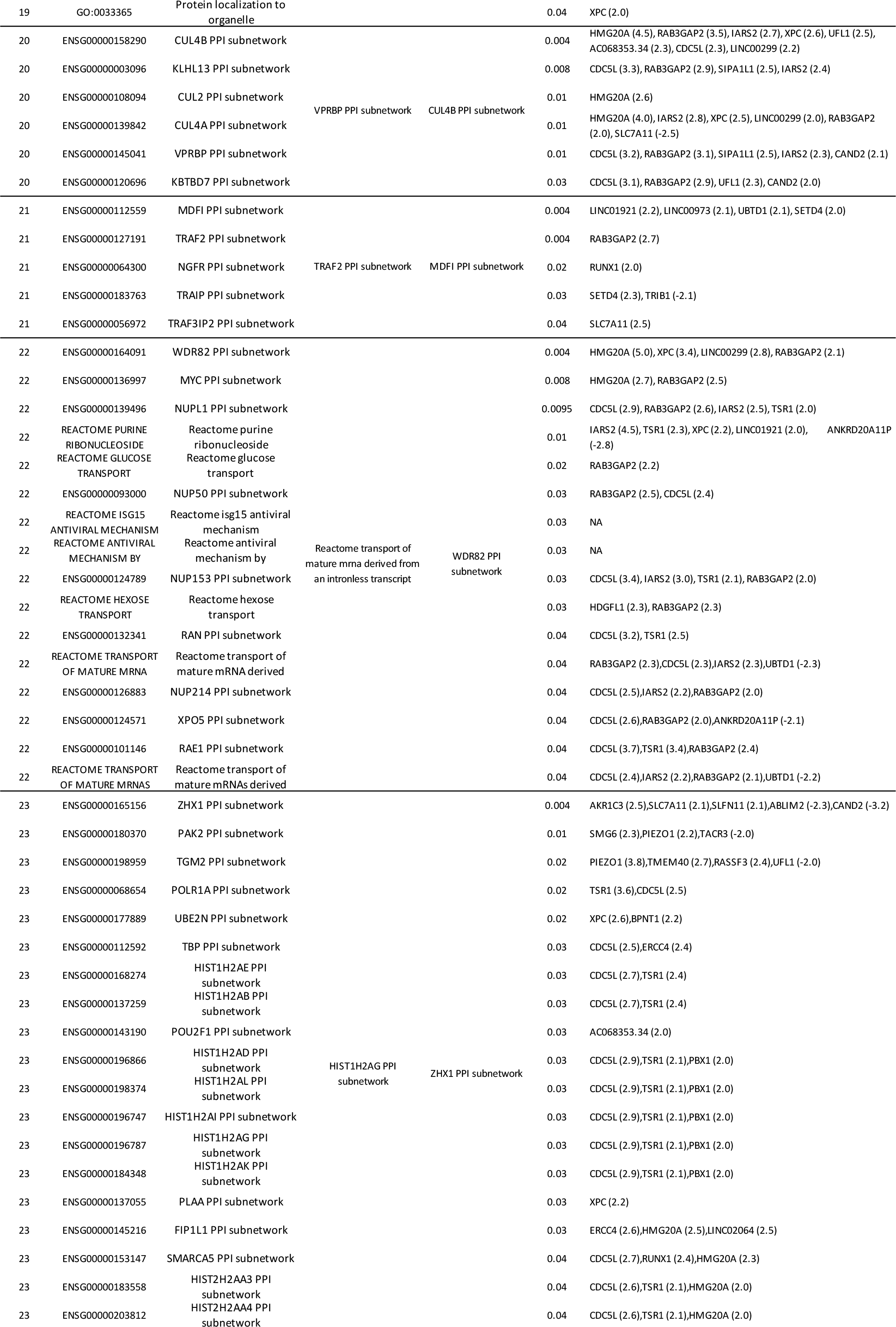

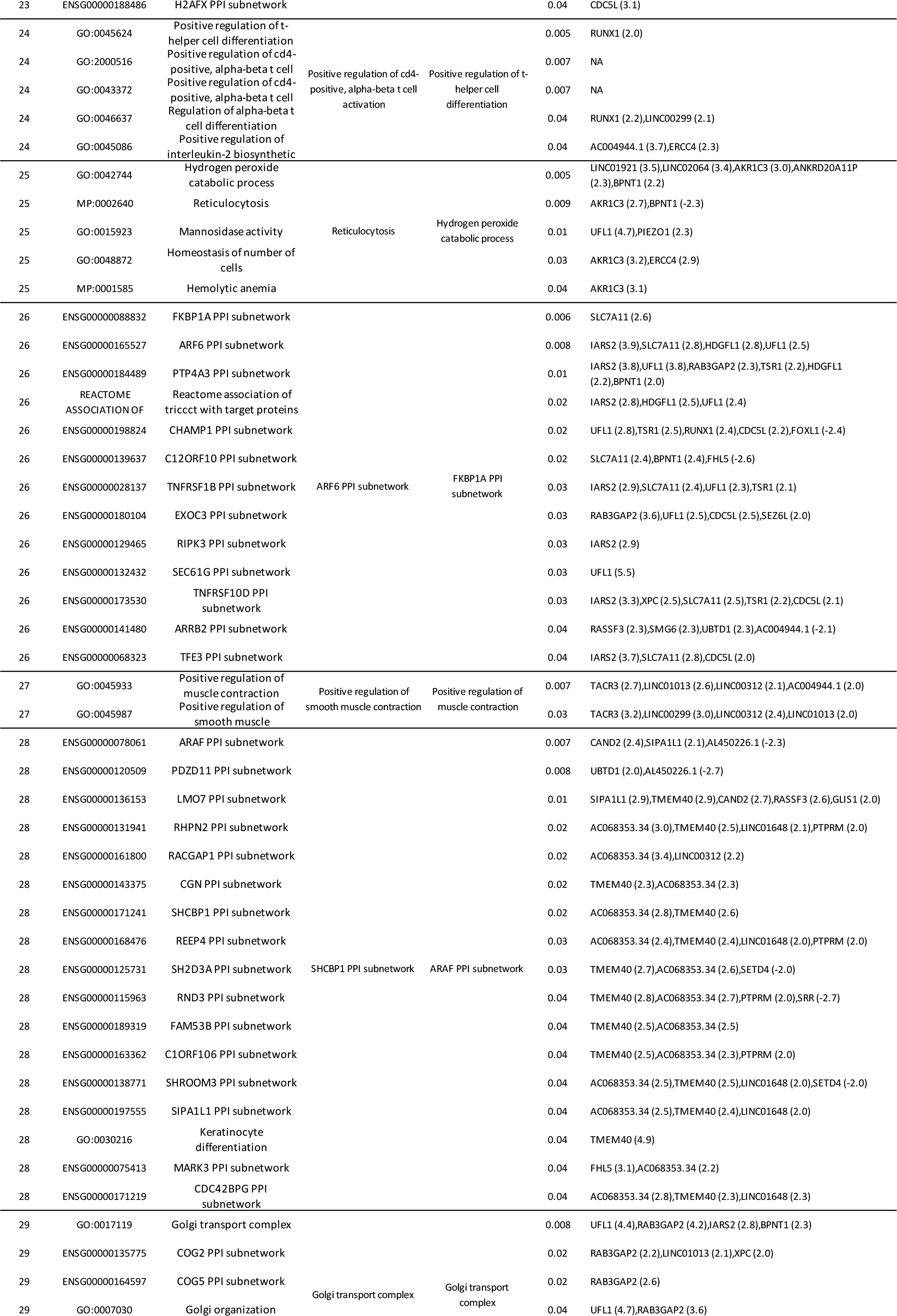

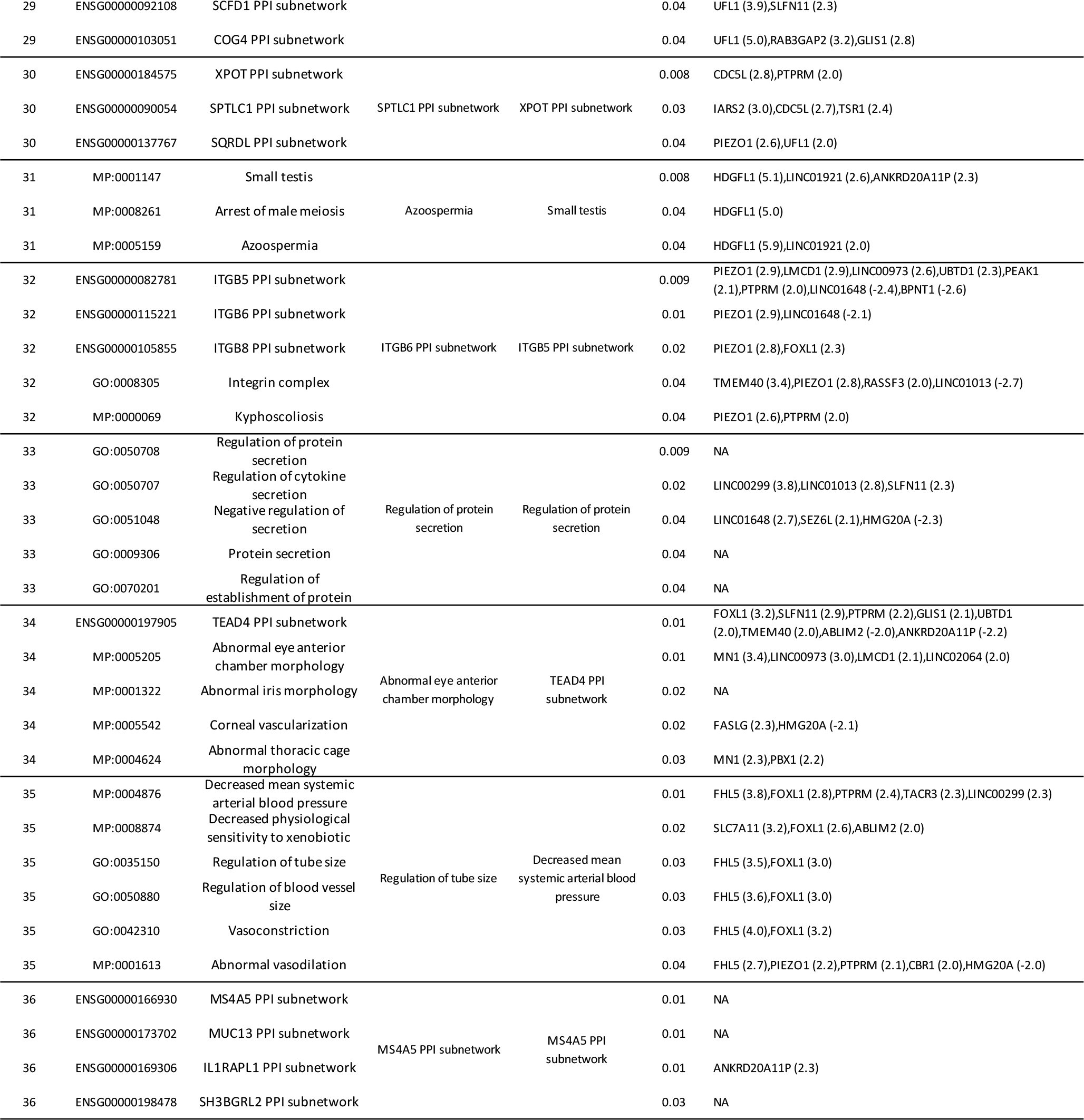

**Supplementary Table 4.**
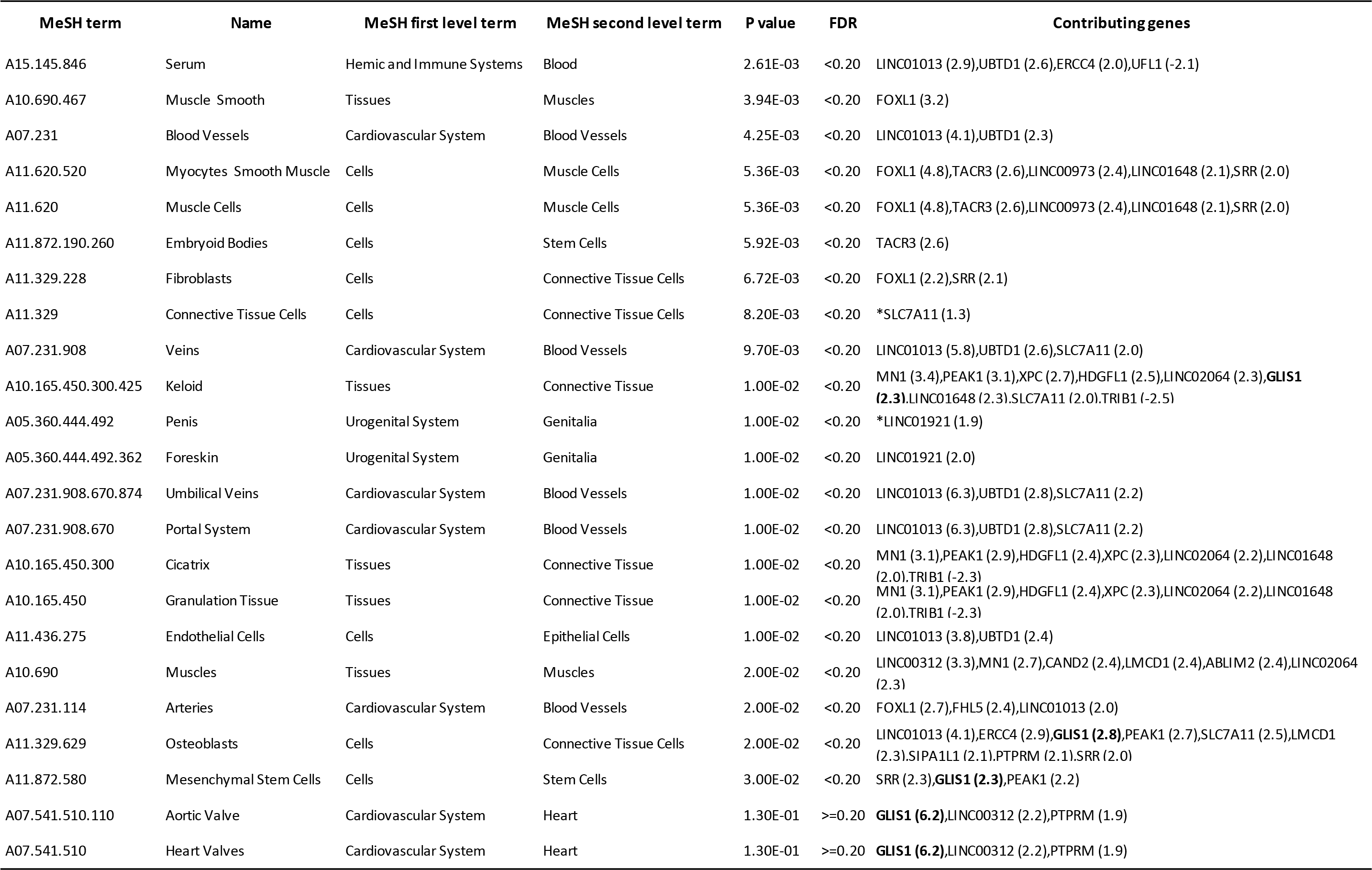
Enrichment of tissue and cell types from DEPICT. Suggestive enrichment (FDR < 0.20) was observed for 21 categories of tissues and/or cell types. Two categories where GILS1 was highly expressed were added. Genes listed were annotated to a given tissue/cell types and are within an associated locus. Each gene has a z-score representing the gene’s likelihood to be highly expressed in the tissue/cell types. Genes displayed are those with a |z-score| > 1.96 that corresponds to a P-value<0.05 as the minimum threshold, or the gene with higher absolute value of z-score (*). MeSH: Medical Subject Headings, FDR: false discovery rate.

**Supplementary Table 5.** The 6 enriched gene sets (FDR < 0.05) where *GLIS1* was found to be the best-ranked gene by i-GSEA4GWAS. Enrichment significance (p-value and FDR) of pathways / gene sets are displayed. Genes where SNPs are associated with MVP (P-value < 0.01 from the GWAS meta-analysis) are listed from the most to the less significant.

| Gene Set Name | P-value | FDR | Significant genes in the pathway/gene set |
| --- | --- | --- | --- |
| GO: Positive Regulation of Transcription DNA Dependent | 0.001 | 0.001 | GLIS1; RUNX1; SMAD3; MKL2; SQSTM1; NUFIP1; NRIP1; PAX8; GLIS3; TBX5; CREB5; ESRRG; MDFIC; RXRA; MYO6; ARNTL |
| GO: Positive Regulation of RNA Metabolic Process | 0.001 | 0.002 | GLIS1; RUNX1; SMAD3; MKL2; SQSTM1; NUFIP1; NRIP1; PAX8; GLIS3; TBX5; CREB5; ESRRG; MDFIC; RXRA; MYO6; ARNTL |
| GO: Positive Regulation of Transcription | 0.001 | 0.003 | GLIS1; RUNX1; SMAD3; HIVEP3; MKL2; IRF4; SQSTM1; NUFIP1; NRIP1; PAX8; GLIS3; TBX5; CREB5; ESRRG; MDFIC; RXRA; NFATC2; MYO6; ARNTL; GLIS2 |
| GO: Positive Regulation of Nucleobasenucleosidenucleotide and Nucleic Acid Metabolic Process | 0.001 | 0.004 | GLIS1; RUNX1; SMAD3; HIVEP3; MKL2; IRF4; SQSTM1; NUFIP1; NRIP1; PAX8; GLIS3; TBX5; CREB5; ESRRG; MDFIC; ABCA1; RXRA; NFATC2; MYO6; ARNTL; GLIS2 |
| GO: Positive Regulation of Transcription From RNA Polymerase II Promoter | 0.001 | 0.009 | GLIS1; RUNX1; SMAD3; MKL2; SQSTM1; NUFIP1; NRIP1; GLIS3; RXRA; MYO6; ARNTL |
| GO: Negative Regulation of Transcription | 0.002 | 0.011 | GLIS1; SMAD3; NFX1; MECP2; NRIP1; TGIF1; GLIS3; BCOR; VDR; TIMELESS; RFX3; ARHGAP35; SUDS3; GLIS2 |
| GO: Negative Regulation of RNA Metabolic Process | 0.023 | 0.089 | GLIS1; SMAD3; NFX1; MECP2; NRIP1; TGIF1; GLIS3; BCOR |
| GO: Specific RNA Polymerase II Transcription Factor Activity | 0.044 | 0.119 | GLIS1; GLIS3; SOX9; RXRA |

**Supplementary Table 6.**
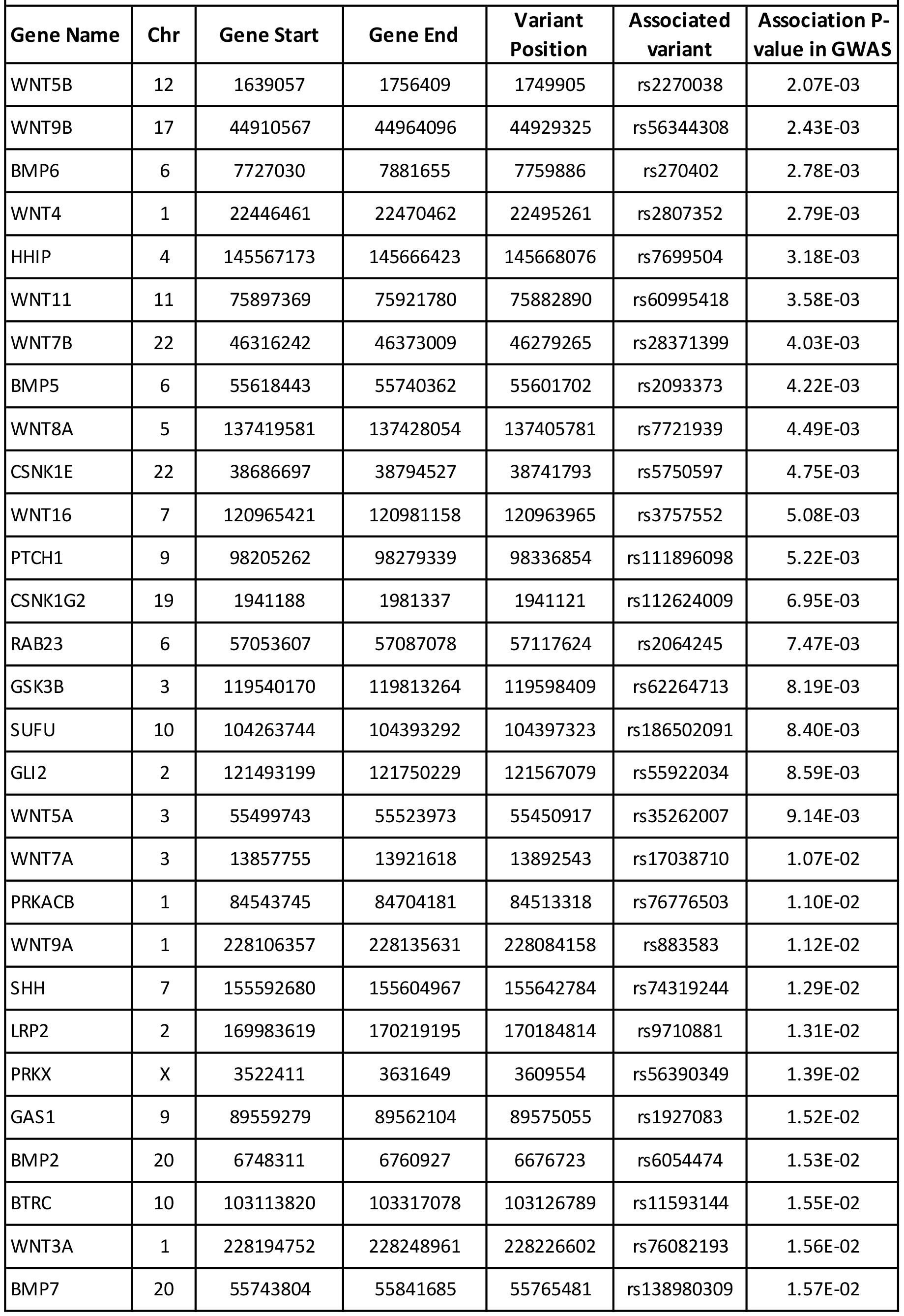
List of 38 significant genes in the Hedgehog signalling pathway (P-value = 0.015) identified by i-GSEA4GWAS. Chr: chromosome. Position from the Hg19 human genome assembly. The rs number and p-value from the GWAS meta-analysis are indicated for the top associated variant in the contributing genes.

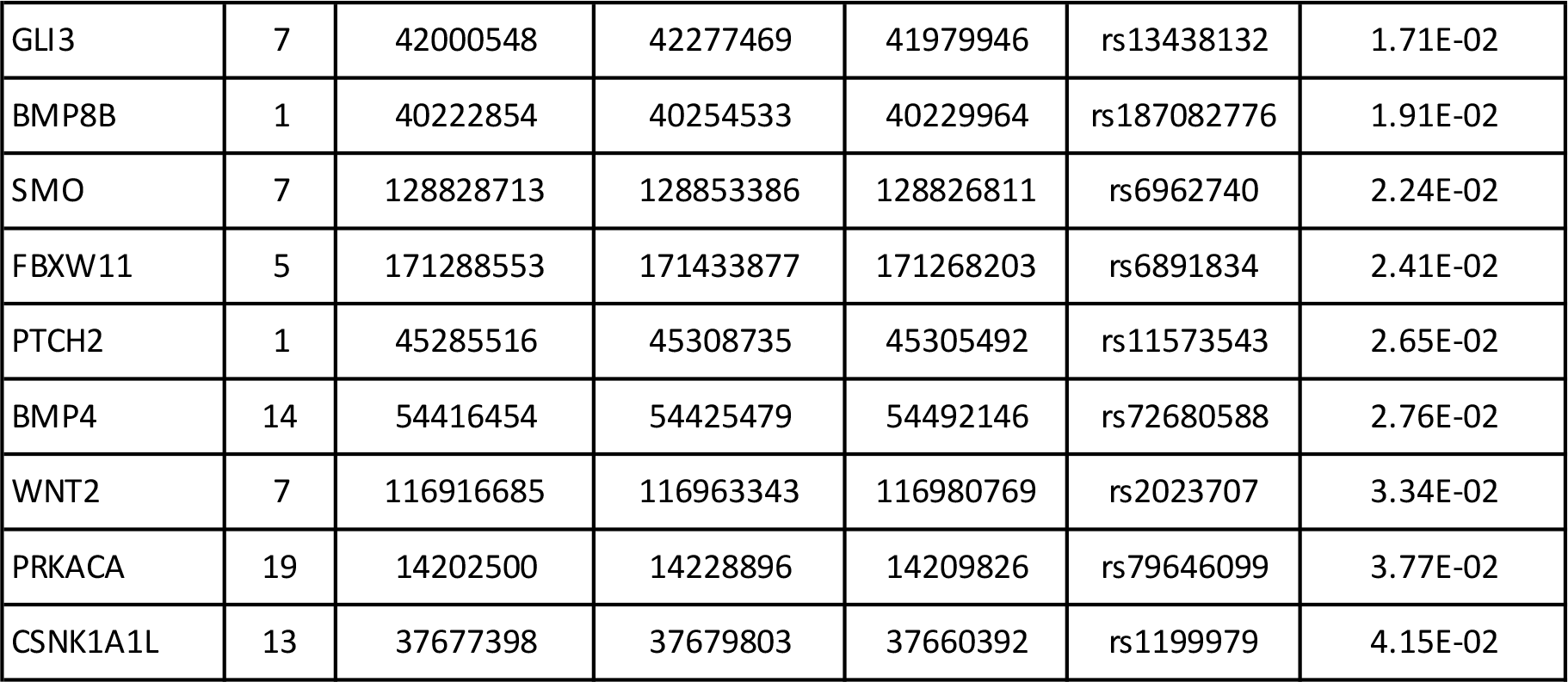

**Supplementary Table 7.** Association of rs1879734 in the discovery meta-analysis and follwo-up in four case control studies

|  | MVP Cases | Controls | OR [95%CI] | P-value |
| --- | --- | --- | --- | --- |
| MVP-Nantes | 489 | 873 | 1.44 [1.18-1.76] | $3.20 \times 10^{-4}$ |
| MVP-France | 953 | 1566 | 1.23 [1.07-1.41] | $3.32 \times 10^{-3}$ |
| **Discovery meta-analysis** | | | | $6.80 \times 10^{-6}$ |
| MGH/FHS | 699 | 5575 | 1.21 [1.05-1.40] | $1.40 \times 10^{-4}$ |
| HEGP-Surgery | 450 | 820 | 1.22 [1.01-1.48] | 0.037 |
| PROMESA | 171 | 282 | 1.06 [0.77-1.46] | 0.88 |
| IUPCQ | 102 | 102 | 0.89 [0.56-1.44] | 0.64 |
| **Global meta-analysis** | 2864 | 9218 | 1.23 [1.14-1.33] | $1.29 \times 10^{-7}$ |
| **Heterogeneity** |  |  |  | 0.41 |

\* N: Numbers; OR: odds ratio; 95%CI: 95% confidence interval
† Alleles are indexed to the forward strand of NCBI Build 37
‡ P-values reported are two-sided and based on an inverse-variance weighted meta-analysis model (fixed effects)
§ The P-value for heterogeneity corresponds to Cochran's Q statistic.

**Supplementary Table 8.**
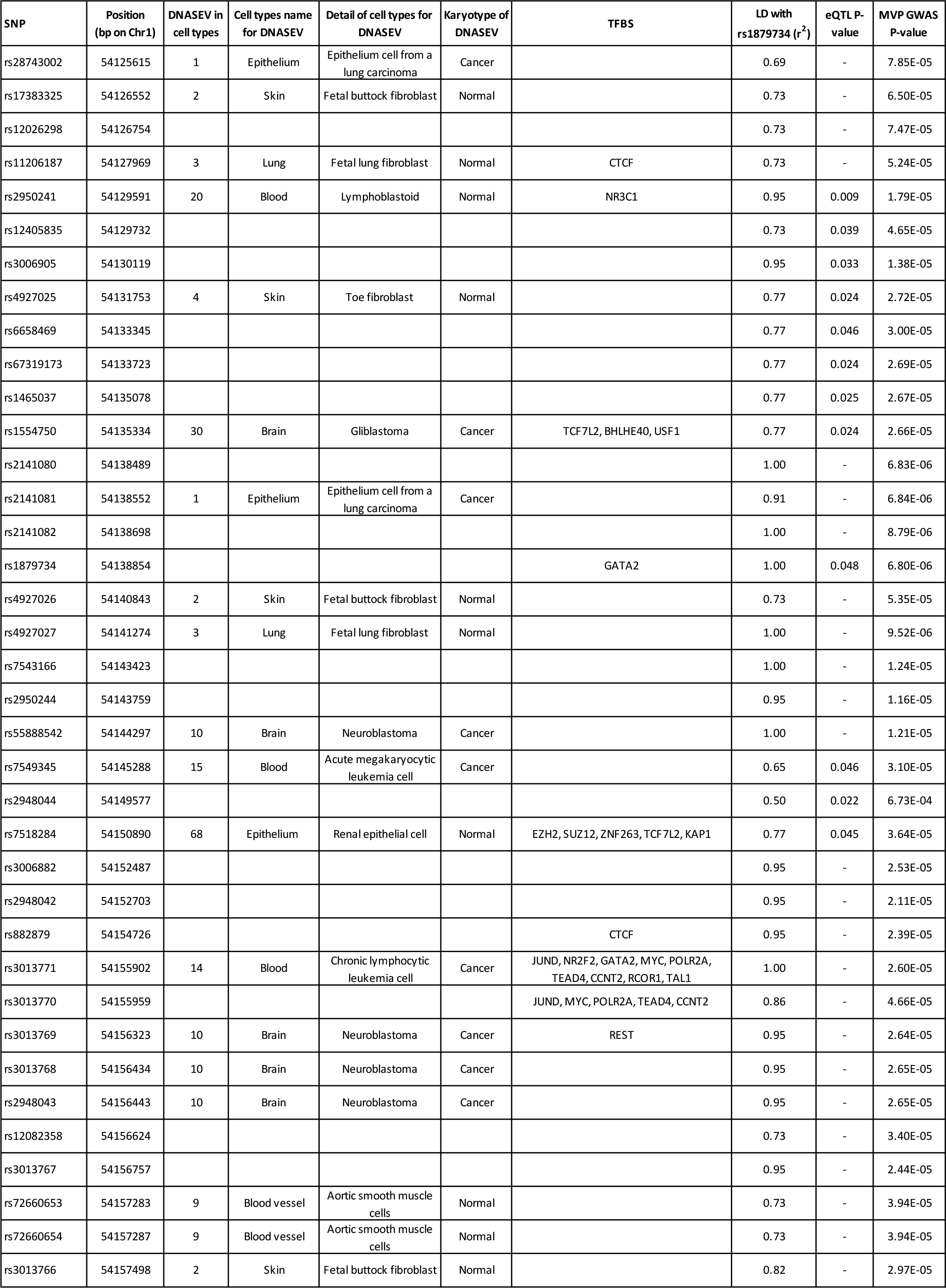
Functional annotation of 111 SNPs at the GLIS1 locus. We provided functional annotations for all nominally associated (P-value < 0.05) SNPs located within ±500Kb and in linkage desequillibrium (r2 > 0.5) with the top associated SNP rs1879743 at the GLIS1 locus. We used 111 SNPs, all were located in intron X as input to Variant Annotation Integrator (VAI), a tool box avialable at the UCSC Genome Browser to annotate SNPs. We provide location in DNase hypersensitive region (DNASEV), cell types and descriptive and the karyotype of cells, in addition to location at transcription factor binding site (TFBS) using ENCODE data. When available, we indicated if SNPs are expression quantitative trait loci (eQTLs) in atrial appendage tissue using GTeX data

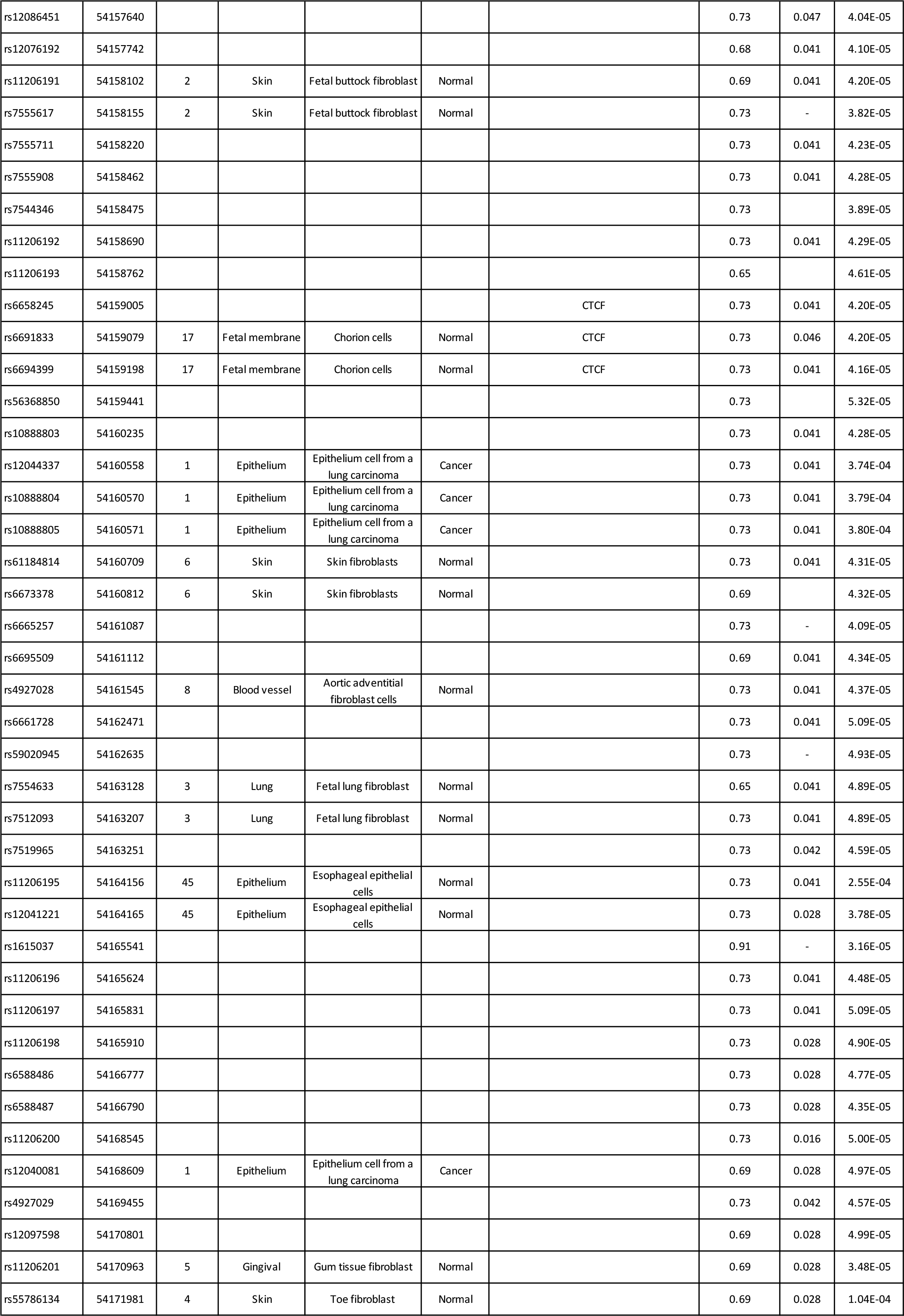

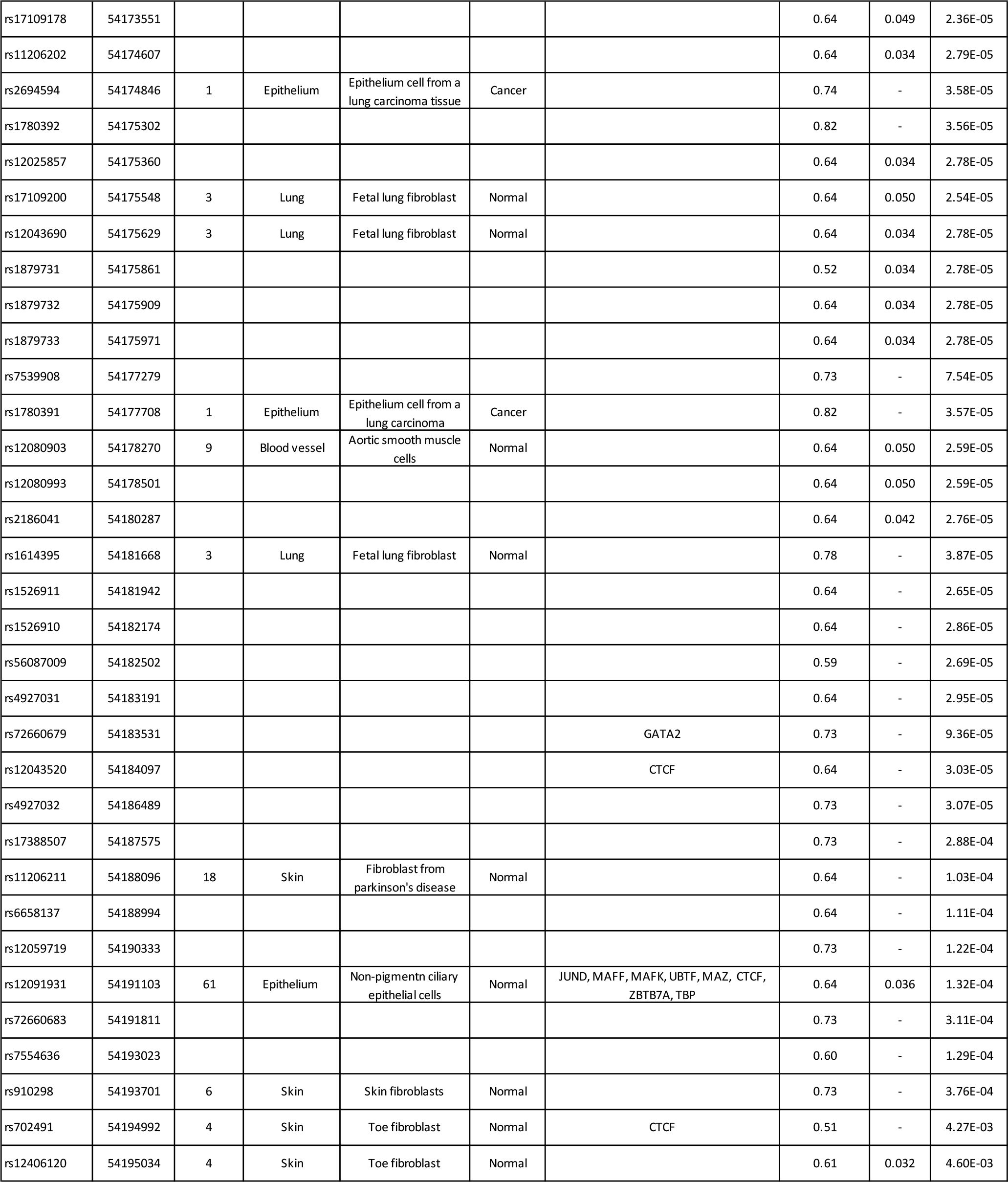

